# Deadenylation rate is not a major determinant of RNA degradation in yeast

**DOI:** 10.1101/2023.01.16.524186

**Authors:** Léna Audebert, Frank Feuerbach, Laurence Decourty, Abdelkader Namane, Emmanuelle Permal, Gwenaël Badis, Cosmin Saveanu

## Abstract

Gene expression and its regulation depend on mRNA degradation. In eukaryotes, degradation is controlled by deadenylation rates, since a short poly(A) tail is considered to be the signal that activates decapping and triggers mRNA degradation. In contrast to this view, we show that global stability of mRNAs can be explained by variations in decapping speed alone. Rapid decapping of unstable mRNAs, for example, allows little time for deadenylation, which explains their longer than average poly(A) tails. As predicted by modeling of RNA degradation kinetics, mRNA stabilization in the absence of decapping led to a decrease in the length of the poly(A) tail, while depletion of deadenylases only increased the tail length. Our results suggest that decapping activation dictates mRNA stability independent of the deadenylation speed.

**One-Sentence Summary:** Unstable mRNAs are characterized by rapid 5’ cap removal, independent of a prior shortening of the poly(A) tail.

## Main Text

Addition of poly(A) tails to RNAs is a major co-transcriptional event during mRNA formation in eukaryotes. Poly(A) tails assist RNA export from the nucleus, its translation and have a protective effect from RNA degradation in the cytoplasm (reviewed in Passmore & Coller, 2021). This protective effect is lost when the poly(A) size decreases to a length of 10-12 nucleotides, a situation that is recognized by the Lsm1-7/Pat1 complex to recruit and activate decapping, committing mRNA to degradation (Tharun *et al*, 2000; Bouveret *et al*, 2000). Thus, the speed of deadenylation, which leads to oligoadenylated species, is widely considered to be a major determinant of RNA instability. This speed is modulated by the activity of the highly conserved Ccr4-Not deadenylase complex (reviewed in Collart, 2016). Specific subunits of the complex, such as Not5, were proposed to couple translation of non-optimal codons with fast mRNA degradation in yeast (Buschauer *et al*, 2020; Webster *et al*, 2018). However, the importance of deadenylation for mRNA degradation in this case and the mechanism leading to decapping activation remain to be demonstrated.

Despite the importance given to deadenylation speed in RNA stability and its regulation, prior deadenylation is not always required for RNA degradation. For example, nonsense-mediated mRNA decay (NMD) is a degradation pathway in which deadenylation does not play a role (Muhlrad & Parker, 1994). In this case, decapping activation is influenced by the frequency at which translating ribosomes encounter a premature termination codon (PTC), as measured by *in vivo* single molecule imaging (Hoek *et al*, 2019). Decapping activation occurs probably at variable rates even for mRNAs that are not targets of NMD, as suggested by kinetic modeling of experimental RNA decay experiments in yeast (Cao & Parker, 2001) or mammalian cells (Eisen *et al*, 2020). These analyses suggest that once an mRNA is deadenylated, its oligoadenylated state is degraded at highly variable speeds, that can vary by a factor of 1000.

How deadenylation rates are causally linked with mRNA degradation remains unclear, in particular when the most stable mRNAs have shorter average poly(A) tails, as shown by large-scale approaches in several organisms, including *C. elegans* (Lima *et al*, 2017) or *A. thaliana* (Jia *et al*, 2022). The correlation between stability of an mRNA and its shorter than average poly(A) tail suggests that deadenylation is not necessarily a causal element in RNA degradation. To clarify the importance of deadenylation speed in mRNA degradation, we performed a comprehensive re-evaluation of RNA degradation models and how they predict large-scale experimental poly(A) tail length and mRNA stability results. We also tested the predictions of these models by inactivation of deadenylation and decapping and measurements of poly(A) tails and mRNA levels on a large-scale. Inefficient deadenylation did not change the degradation rate of reporter RNAs, despite the presence of extended poly(A) tails. Our experimental and computational results suggest that deadenylation speed is not a limiting step in yeast. The obtained results are compatible with large-scale data and correspond to predictions of a model in which deadenylation speed and decapping activation are independent events.

## Results

We became interested in the relationship between poly(A) tail size and mRNA degradation when analyzing the proteins associated to NMD complexes, in particular Upf1 (Dehecq *et al*, 2018). These complexes contain both Pab1, a protein that efficiently binds long poly(A) tails (Schäfer *et al*, 2019) and Lsm1, component of the Lsm1-7/Pat1 complex, which binds preferentially to oligoadenylated RNA (Chowdhury *et al*, 2014). We characterized the protein composition of complexes associated with Pab1 (**Fig. 1A**), Lsm1, Pat1, Lsm7 and Dhh1(**Fig. S1A to D**) and found that, in every case, Upf1 was among the enriched proteins (**Data Set S1**). The results were specific, since, for example, Lsm2, present in most of the purified complexes, was absent from Pab1-TAP (**Fig. S1E**). In support to the hypothesis that the majority of observed interactions with Ufp1 were mediated by RNA, the association of Pab1-HA with Upf1 was sensitive to a nuclease treatment (**Fig. 1B**).

**Fig. 1.**
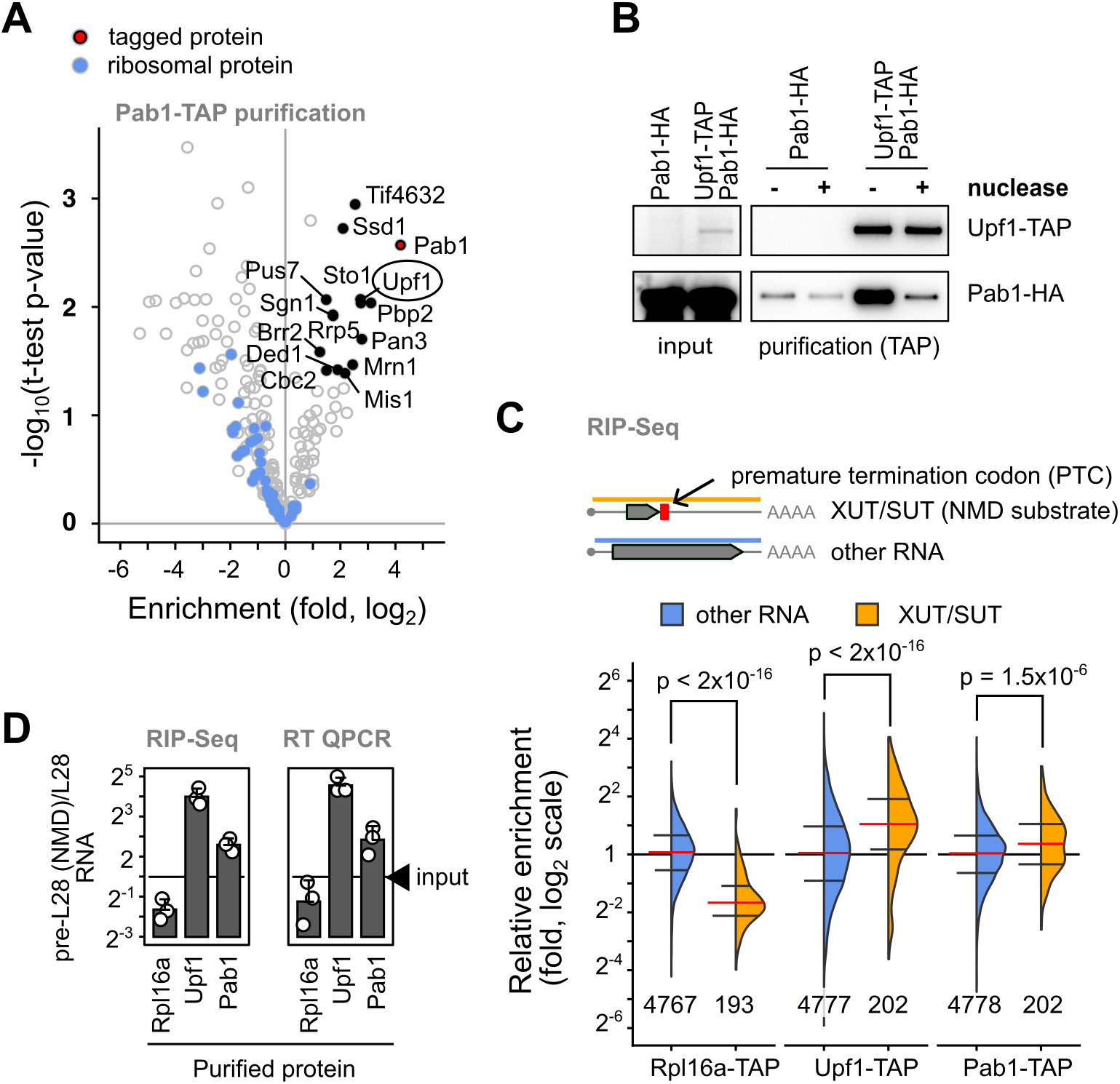
NMD substrates have long poly(A) tails and are associated with Pab1. **(A)** Enrichment of proteins in association with Pab1-TAP was estimated by quantitative mass spectrometry for identified proteins in comparison with their abundance in a total protein extract. Labeled proteins correspond to those for which at least three independent measures were available, proteins were enriched by a factor higher than 2 and were robustly detected, as judged by a *p-value* lower than 0.05 for a t-test (null hypothesis). Blue dots indicate ribosomal proteins. **(B)** The interaction between Pab1 and Upf1 was tested by co-purification of Pab1-HA with Upf1-TAP in the presence or absence of micrococcal nuclease treatment. **(C)** Relative enrichment of NMD substrates, annotated as “XUT/SUT”, in association with Rpl16a-TAP, Upf1-TAP and Pab1-TAP, showed the preference of Upf1 and Pab1 to this unstable RNA population (orange), in comparison with other cellular RNA (blue). The red horizontal line indicates the median of the enrichment values, with the first and third quartiles indicated by horizontal black lines. The indicated p-values correspond to a Wilcoxon 15 rank sum test with continuity correction, N is the number of measured transcripts in each category. **(D)** Validation of RIP-Seq experiments by independent RT-qPCR tests of the relative enrichment of a known NMD substrate, the unspliced pre-mRNA for RPL28 and its non-NMD equivalent, the spliced mRNA. The ratio between the two forms of RNA were compared to the ratio in the total RNA sample (input).

To explore the potentially different populations of RNA molecules present in Lsm1-7/Pat1 and Pab1-associated mRNPs together with Upf1, we sequenced the RNA co-purified with Lsm1-TAP, Pab1-TAP and Upf1-TAP. A purification of Rpl16a-TAP was done in parallel, as it allows an estimation of the levels of RNA bound to ribosomes (Halbeisen *et al*, 2009). The enrichment profiles for RNA with the tagged proteins, were, with the exception of the Lsm1-TAP situation, different from a control purification performed with a strain that did not express any tagged protein. We thus focused on the results obtained with purifications of Upf1, Rpl16a and Pab1.

To validate the obtained results, we analyzed first a sub-category of yeast RNA, the Xrn1-dependent unstable transcripts (XUT, van Dijk *et al*, 2011) and the stable unannotated transcripts (SUT, Xu *et al*, 2009) that are known to be unstable and frequently targets of the NMD pathway (Malabat *et al*, 2015). As a positive control, we observed a significant enrichment of XUT/SUT transcripts in the Upf1 purification, while XUT/SUT RNAs were strongly depleted from the ribosome associated fraction (**Fig. 1C**). This depletion was expected, since XUT/SUT only contain spurious short coding sequences and, like other NMD substrates (Heyer & Moore, 2016), are probably at most associated with a single translating ribosome. Surprisingly, XUT/SUT were enriched in the Pab1 associated fraction. To confirm this observation, we analyzed pre-mRNAs for ribosomal protein genes as a different type of NMD substrate and compared intron-containing transcripts with the spliced ones. As expected, and similar to the XUT/SUT situation, pre-mRNAs were significantly depleted from the ribosome associated fraction, strongly enriched in association with Upf1, but also enriched in the Pab1 bound RNA population (**Fig. S2A**, example in **Fig. S2B**, data in **Data Set S2**). The relative enrichment of the pre-L28 NMD sensitive transcript in Upf1-TAP and Pab1-TAP purifications in comparison with the spliced RPL28 mRNA was confirmed by RT-qPCR (**Fig. 1D**).

We found Pab1 to be associated with unstable transcripts that are also bound by NMD factors, which probably reflects their fast degradation through a deadenylation-independent mechanism. Since rapid degradation does not allow deadenylation to occur, long poly(A) tails of NMD substrates are a marker of RNA instability. Interestingly, and independent of their NMD status, longer poly(A) tail transcripts were on average enriched in the Pab1-associated fraction in comparison with shorter poly(A) mRNA (**Fig. S2C**). We conclude that Pab1 preferentially binds long poly(A) tail RNAs and that the association with Pab1 can be a marker of RNA instability.

The unexpected finding that Pab1 binds unstable RNAs with relatively long poly(A) tails, led us to explore the relationship between RNA half life and poly(A) tail length from published large-scale data sets. Poly(A) tail length was strongly inversely correlated with RNA stability (**Fig. 2A, 2B, S3A, B**) as previously reported (Chan *et al*, 2018; Lima *et al*, 2017). The observed correlations were independent on the method used for measuring RNA half-life or poly(A) tail length. If deadenylation speed dictates RNA stability, changes in deadenylation activity should be reflected in predictable changes in poly(A) tail lengths for both stable and unstable RNA. To test this hypothesis, we analyzed the recently published changes in poly(A) tail length in strains lacking Pan2 and Ccr4 deadenylases (Tudek *et al*, 2021). Most stable 25% RNAs and most unstable 25% RNAs showed opposite relative changes in poly(A) tail length (**Fig. S3C**). Slowing down deadenylation led to variations in poly(A) tail length that were not clearly linked with RNA stability, leading to the question of the impact of deadenylation on RNA degradation in yeast.

**Fig. 2.**
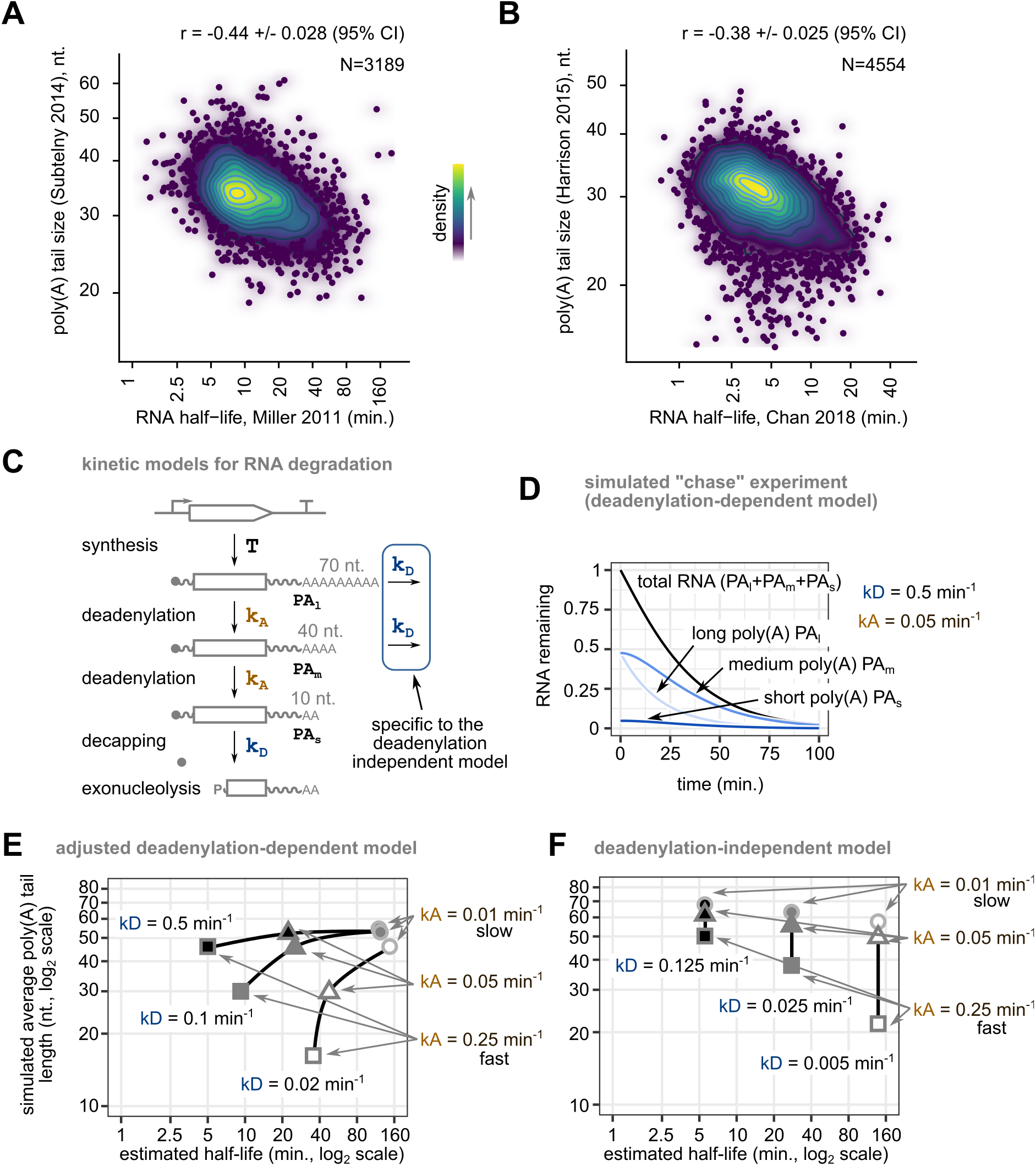
The presence of long poly(A) tails on unstable mRNAs can be explained by two RNA degradation mechanisms. **(A)** Average poly(A) tail length shows a negative correlation with RNA half-life. Published poly(A) tail size data (log2, Subtelny *et al*, 2014), were represented on the vertical axis as a function of estimated RNA half-life, horizontal axis (log2, Miller *et al*, 2011). The Pearson product moment correlation coefficient and its 95% confidence interval are displayed on the graph. **(B)** Similar to panel (A), but with data for poly(A) size and RNA half-life from other publications (Harrison *et al*, 2015; Chan *et al*, 2018). **(C)** Depiction of two models for RNA degradation, one in which deadenylation is required to induce decapping, and another in which decapping can occur on RNAs with any size of poly(A) tail. k_A_ is a kinetic constant for deadenylation, k_D_ represents a decapping rate constant and “T” indicates all the processes leading to the generation of cytoplasmic mRNA. **(D)** Example of simulated RNA degradation curve for the deadenylation-dependent kinetic model, starting from a steady-state situation described by a decapping constant of 0.5 min^-1^ and a deadenylation constant of 0.05 min^-1^. The relative variation of the amounts of the three forms of RNA is indicated. **(E)** Examples of simulated half-life values and average poly(A) tail length for the deadenylation-dependent model for RNAs with three values of k_D_ (low, white, intermediate, gray, and high, black) in combination with three values of k_A_ (low, circles, intermediate, triangles, and high, squares). Values of poly(A) for the three species were arbitrarily set to 70, 40 and 10 nucleotides to calculate averages. Both axes are logarithmic. **(F)** Similar to (E), but for the deadenylation-independent kinetic model. The k_D_ values were adjusted to obtain half-life 10 and poly(A) size values compatible with published results.

Previous analyses of RNA degradation assumed that deadenylation is the critical step in RNA degradation, with rapid degradation of RNA once it reaches a critical size of the poly(A) tail, of about 10 to 12 nucleotides (Muhlrad *et al*, 1994; Cao & Parker, 2001). We wondered how well this deadenylation-dependent model for RNA degradation fits the observed short poly(A) tail length of most stable mRNAs. To this end, we used a kinetic model for RNA degradation inspired from previous work (Cao & Parker, 2001). Three poly(A) species were considered for each RNA, starting with long poly(A) tail RNA (PA_l_), which is transformed to intermediate (PA_m_) and short tail RNA (PA_s_) at a speed proportional with a constant related with deadenylation, k_A_ (**Fig. 2C**). Oligoadenylated RNA is finally degraded with a first order constant k_D_. The functions that express the concentration of each RNA species as a function of time were obtained by solving the set of differential equations for the presented model (**Fig. S3E**). A deadenylation-independent model was also considered, in which degradation can also occur for the longer poly(A) tail species (**Fig. 2C, Fig. S3F** for the corresponding equations). These models were equally employed within *Tellurium*, a modeling and simulation software package (Choi *et al*, 2018) using the representation depicted in **Fig. S3D**. The computed steady-state levels for RNAs of different poly(A) tail length were used as a starting point in simulated “chase” experiments, to calculate both the variation in average poly(A) tails and amount of remaining RNA (**Fig. 2D**, for an example).

To understand the impact of deadenylation and decapping speeds on simulated half-life and poly(A) tail distribution, we varied both k_A_ and k_D_ values. For a constant high k_D_ of 0.5 min^-1^, changing kA values by a factor of 25 led, for example, to a shift in the half-life for the simulated RNA from 5 to 120 minutes (**Fig. 2E, upper line**). With arbitrary values for the three poly(A) species of 70 (PA_l_), 40 (PA_m_) and 10 (PA_s_) nucleotides (**Fig. 2C**), we computed average sizes of the poly(A) tails at steady-state in each condition. The obtained values ranged from 46 nucleotides for the fastest decaying RNA to 55 nucleotides for the slowest ones (**Fig. 2E, upper line**). This result is in opposition with the experimentally observed long poly(A) tails for unstable mRNAs (**Fig. 2A, B**). To solve this conundrum we tested the effect of changing the degradation rate of oligoadenylated RNA, k_D_. As expected, decreasing k_D_ by a factor of 5, from 0.5 min^-1^ to 0.1 min^-1^ led to an increase in the RNA half-life from 5 to 9 minutes and a decrease in the average length of poly(A) tails at steady-state from 46 to 30 nucleotides (**Fig. 2E**). Further decreasing k_D_ led to an increase in RNA half-life from 9 to 35 minutes, and a decrease in average poly(A) tail size of RNAs from 30 to 16 nucleotides (**Fig. 2E**). Thus, k_D_ changes, rather than deadenylation speed might be responsible for the observed correlation between poly(A) size and RNA instability (**Fig. 2A, B**).

An alternative explanation for the presence of long poly(A) tails for unstable mRNA is that mRNA degradation follows a deadenylation-independent model. Such a model involves the addition of just two additional degradation steps (**Fig. 2C**). Increasing k_D_ values in this model led to shorter RNA half-lives, with a range of average poly(A) tails length that was dependent on the simulated k_A_ (**Fig. 2F**).

Thus, the inverse correlation between average poly(A) tail length and RNA half life (**Fig. 2A, B**) can be explained by the classical deadenylation-dependent model only when the limiting step for RNA degradation is not deadenylation speed (k_A_) but rather the degradation of the oligoadenylated RNA species (k_D_). Alternatively, the observed anti-correlation can also be explained by a model in which decapping can occur even on long poly(A) tail RNA (**Fig. 2C**), as it happens for NMD substrates. The two modes of RNA degradation are not mutually exclusive but can be experimentally tested by modulating deadenylation or decapping.

Slowing down deadenylation is expected to lead to the rapid accumulation of unstable RNAs in the classical model and to have no impact in deadenylation-independent RNA degradation (**Fig. S4A, B**). Decapping inhibition should lead to a rapid increase in the levels of unstable RNA in both models (**Fig. S4C, D**). To test these predictions, we set up conditions in which the major yeast deadenylases Ccr4 and Pop2/Caf1, or the decapping enzyme Dcp2, were rapidly depleted. The system (**Fig. S5A**) is based on the fusion of the protein of interest with an auxin-inducible degron (AID) domain recognized by the auxin-sensitive *O. sativa* TIR1 (Nishimura *et al*, 2009), whose expression is induced by a β-estradiol treatment. Addition of β-estradiol and indole-3-acetic acid (IAA) auxin triggered proteasome-degradation of proteins, as seen by the rapid decrease in the levels of Ccr4, Pop2 or both proteins (**Fig. 3A**). This decrease in protein levels was accompanied by a slow-growth phenotype, only visible under induction conditions (**Fig. S5B**).

**Fig. 3.**
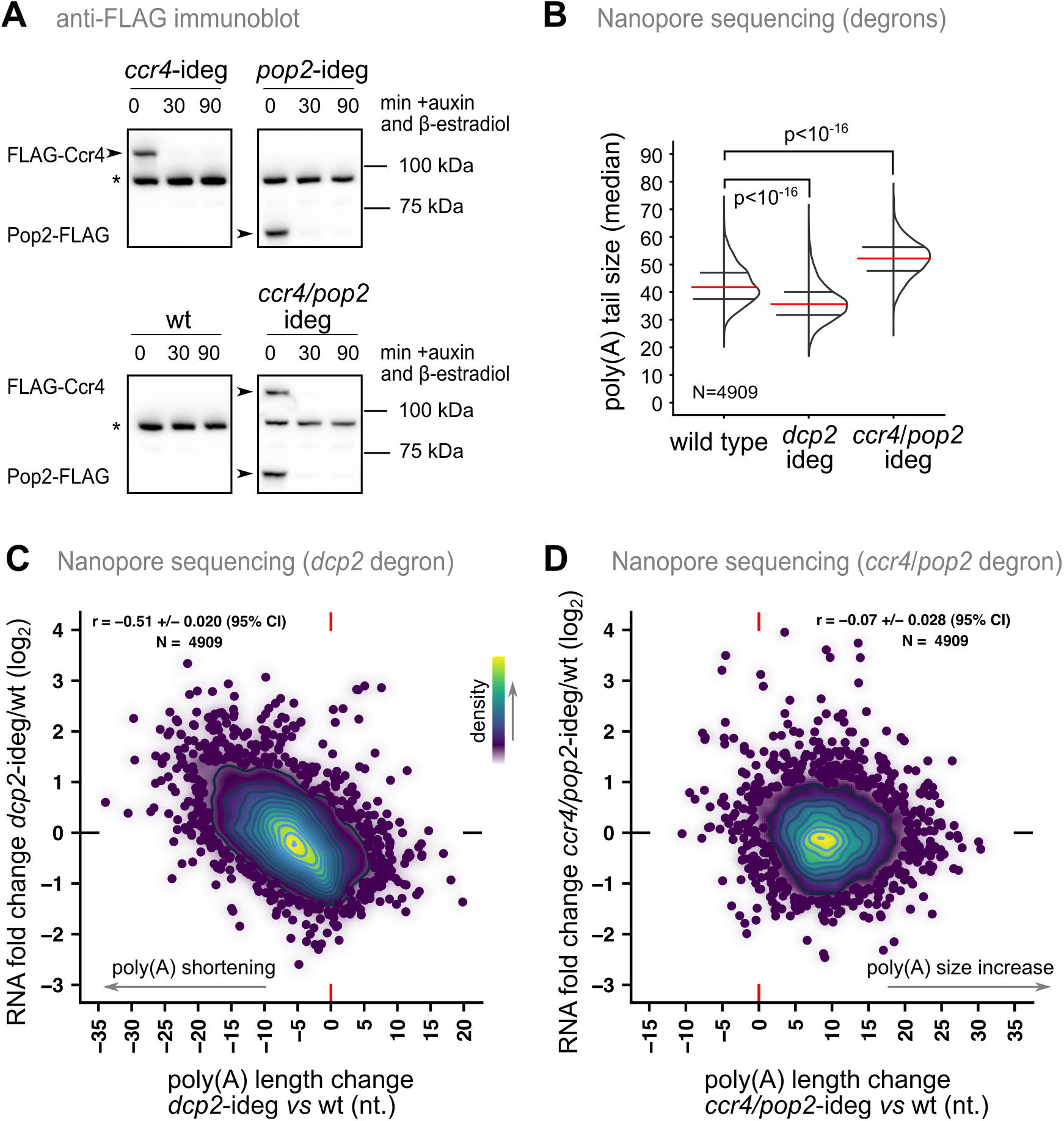
Poly(A) tail changes when decapping or deadenylation are inactivated suggest a deadenylation-independent model for RNA degradation. **(A)** Immunoblot estimation of the decrease in the levels of deadenylases targeted by the inducible degron system. Protein degradation was induced by addition of β-estradiol (1 μM) and IAA (100 μM) and total protein extracts were tested at the indicated time points with anti-FLAG antibodies. The asterisk indicates a yeast protein that is detected by the antibodies in all samples. **(B)** Comparison of the global distribution of median poly(A) tails for mRNAs as detected by Nanopore sequencing in strains depleted for Dcp2 or Ccr4 and Pop2 by treatment with β-estradiol and IAA for 1 hour in liquid medium. The results represent the average of two independent experiments. Red lines correspond to medians of the distributions, with black lines indicating the first and the third quartiles. Indicated *p-values* correspond to a paired Wilcoxon signed rank test with continuity correction for the null hypothesis. **(C)** Relationship between the measured changes in poly(A) tail length in Dcp2-depleted cells and the increase in the levels of the corresponding mRNAs, compared with a wild type strain. The Pearson product moment correlation coefficient and its 95% confidence interval are displayed on the graph. **(D)** Similar to (C) for the strain depleted for the Ccr4 and Pop4 deadenylases.

We used Nanopore sequencing to estimate both the changes in poly(A) tail length and RNA levels (Tudek *et al*, 2021) in strains depleted for the two deadenylases of the Ccr4-Not complex and for Dcp2. Depletion of these proteins led to global changes in poly(A) tail length. As expected, mRNA poly(A) tails became globally longer when both deadenylases were depleted, with an average shift from 42.5 to 52 nucleotides (**Fig. 3B**). Depletion of Dcp2 had the opposite effect, with a decrease of the average poly(A) tail size to 36.3 nucleotides. This shift was expected if Dcp2 depletion stabilizes mRNAs, and leaves more time for deadenylation to occur. As we were interested in the impact of poly(A) tail changes on RNA levels we looked at how transcript poly(A) tail length changed in the degron strains in relation with RNA amounts (**Data Set S3**). We found that an increase in mRNA levels was accompanied by a decrease in the size of the poly(A) tails when Dcp2 was depleted (**Fig. 3C**). Such correlated changes were not observed in the strains depleted for the Ccr4 and Pop2 deadenylases (**Fig. 3D**), even if poly(A) tail size was clearly and globally increased. Perturbation of deadenylation or decapping led to results that are compatible with a deadenylation-independent degradation of unstable RNAs in yeast.

Our results indicated that deadenylation-independent degradation affects a sizable fraction of yeast mRNAs. Since NMD could be responsible for part of this observed effect, we used a reporter system for RNAs that are not sensitive to NMD to estimate the dynamics of RNA levels and of the poly(A) tail when decapping or deadenylation were inactivated. To compare RNAs with different stabilities, we took advantage of the previous observation that coding sequence bias affects RNA degradation (Presnyak *et al*, 2015; Herrick *et al*, 1990) and built two reporter RNAs with HIS3 coding sequences that were either optimal (OPT-HIS3) or suboptimal (non-OPT-HIS3) in terms of codon usage. A tetOFF system was used to block reporter synthesis following the addition of doxycycline (**Fig. 4A**). Half-life estimates for the two reporter mRNAs were of 16 minutes for OPT-HIS3 and 8 minutes for non-OPT-HIS3 (**Fig. 4B**). The reporters, and in particular non-OPT-HIS3, were stabilized by the addition of the translation inhibitor cycloheximide (**Fig. S6A**) and were not affected by NMD inhibition (**Fig. S6B, C**).

**Fig. 4.**
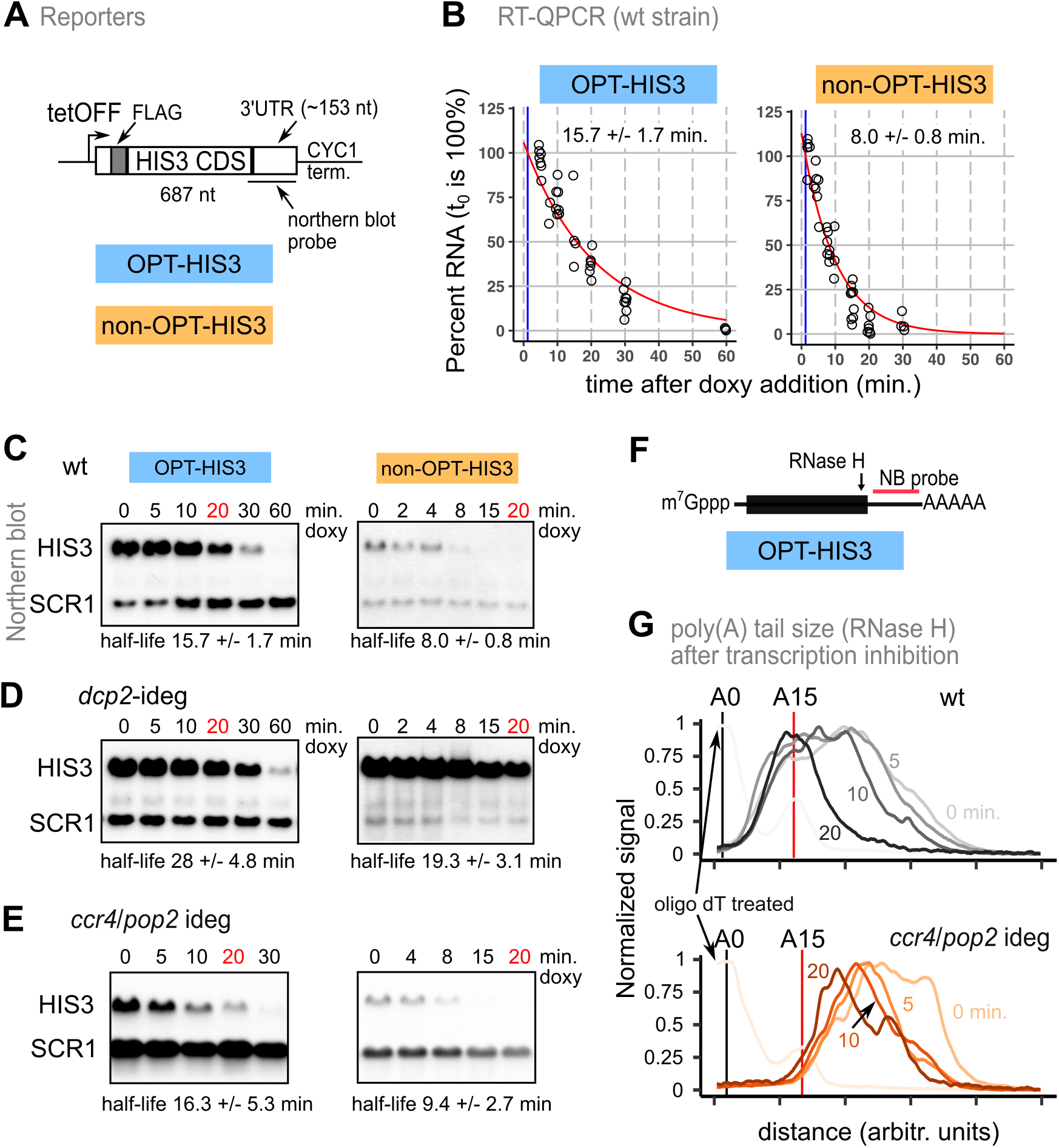
Deadenylation speed does not affect half-life of reporter mRNAs. **(A)** Reporter mRNA controlled by a tetOFF system were used for half-life estimates. **(B)** Estimation of degradation rates for reporters was based on RT-qPCR experiments performed after addition of doxycycline. Estimated mRNA half-life values and 95% confidence intervals are represented. **(C)** Northern blot visualization of degradation for reporter mRNA in comparison with the stable SCR1 RNA, as control. The indicated half-life values correspond to quantitative experiments done by RT-qPCR. **(D)** Similar to (C) but following depletion of Dcp2. **(E)** Similar to (C) but following depletion of deadenylases Ccr4 and Pop2. (F) Estimation of the poly(A) size for reporter RNA was done after RNAse H digestion using northern blots. **(G)** Profiles of poly(A) tails for OPT-HIS3 reporter mRNA at different time points after transcription inhibition in a wild type strain (upper panel) or following depletion of Ccr4 and Pop2. The corresponding northern blot images are presented in **Fig. S8A**.

Next, we tested reporter mRNA levels at different time points after addition of doxycyline in a strain depleted for Dcp2. The half-life of both reporter mRNA doubled (**Fig. 4C, D** and **Fig. S7A**) in these conditions. Intriguingly, the measured half life of the non-OPT-HIS3 and OPT-HIS3 was not affected in degron strains after depletion of Ccr4, Pop2 or both these deadenylases (**Fig. 4E, Fig. S6D, E** and **Fig. S7B, C, D**). These results were correlated with the lack of global mRNA level changes despite the upshift of poly(A) tail lengths measured by Nanopore sequencing (**Fig. 3B, D**). To better understand the impact of blocking deadenylation on reporters poly(A) tail and degradation, we tested both their steady-state and deadenylation dynamics by using RNAse H digestion and Northern blotting (**Fig. 4F, G, Fig. S8A-D**). Depletion of deadenylases, either alone or combined, drastically changed the steady-state distribution of poly(A) tails for both reporters. The most striking change was the decrease in the signal corresponding to relatively short poly(A) tails following deadenylases depletion (**Fig. S8A, B**). At steady-state, poly(A) tails encompassed a broad range, from less than 10 to more than 80 A. However, when deadenylases were depleted, in isolation or at the same time, the signal corresponding to short poly(A) tails was strongly reduced. While short poly(A) tails could not be visible for the reporter RNAs in cells depleted for Ccr4, they could be partially detected in the Pop2 depleted cells (**Fig. S8B-D**). These differences were comparable with those observed for total RNA poly(A) tail dynamics after transcription shut-off, where CCR4 deletion led to a loss of deadenylated species shorter than 20-23 nucleotides (Webster *et al*, 2018).

In conclusion, changes in poly(A) tail length of reporter RNAs had no impact on RNA half-life. Together with the distribution of poly(A) tails of stable and unstable mRNAs, the RNA degradation modeling results and the Nanopore sequencing data, these observations suggest that deadenylation-independent RNA degradation is a more prevalent mechanism than previously thought. An uncoupling of decapping from deadenylation is thus probable not only during meiosis, as recently proposed (Wiener et al, 2021), but during mitotic growth as well. Considering decapping and deadenylation as independent processes explain previously published results and reinforces the idea that “deadenylation is not the rate-limiting event in mRNA degradation” 5 (Herrick et al, 1990). Since the steady-state correlation between long poly(A) tail length and mRNA instability seem to be broadly distributed among various eukaryotes (Lima et al, 2017; Jia et al, 2022; Legnini et al, 2019), our conclusions probably apply to multiple species. A re-evaluation of the deadenylation-dependent and deadenylation-independent models for RNA degradation could help in better understanding the complex relationship between the poly(A) tail, translation and 10 RNA stability. The results presented here should encourage the use of alternative models for RNA degradation in the interpretation of large-scale results to allow the identification of the responsible molecular mechanisms.

## Supporting information

Data Set S3

Data Set S2

Data Set S1

## Acknowledgments

We thank our colleagues of the Genetics of Macromolecular Interactions laboratory for fruitful discussions. We thank Mariette Matondo and Julia Chamot-Rooke for providing access to the Orbitrap Velos mass spectrometer at the Proteomics facility of the Institut Pasteur. We thank Laurence Ma (Biomics Platform, C2RT, Institut Pasteur, Paris) for assistance in DNA sequencing. We thank G. Bushkin, J. Coller, D. Challal and D. Libri for sharing reagents and strains.

## Funding

French ANR grant ANR-18-CE11-0003-04 (CS)

French Ministère de l’enseignement supérieur et de la recherche PhD grant (LA)

Fondation ARC pour la recherche sur le cancer (LA)

## Author contributions

Conceptualization: LA, FF and CS

Experimental work: LA, FF, LD, AN and GB

Data analysis: LA, FF, AN, LD, EP, GB and CS

Data visualization: LA, AN, GB and CS

Manuscript writing: LA and CS

Manuscript editing: all authors

Funding acquisition: LA and CS.

## Data and materials availability

The accession number for the sequencing data reported in this paper is GSE160642 (GEO, Gene Expression Omnibus, NCBI). The MS proteomics data that support the findings of this study have been deposited in the ProteomeXchange repository with the dataset identifier PXD028008. Nanopore sequencing results were deposited in GEO, with accession number GSE211782.

## Supplementary Materials

Materials and Methods

Figs. S1 to S8

Data Set S1 to S3

## Supplementary materials

– Materials and Methods
– Strains, oligonucleotides, plasmids
– Supplementary figures (S1 to S8)

## Materials and Methods

### Yeast and bacterial strains

*Saccharomyces cerevisiae* strains were derived from BY4741 (Mat a) and BY4742 (Mat α) strains. The full list of used strains is provided in the Supplementary material. Strain GBy68 (Bushkin *et al*, 2019) was kindly provided by G. Bushkin, Whitehead Institute for Biomedical Research, Cambridge, MA, USA. An alpha haploid spore derived from this strain, with the *rme1*-Δ, ins-308A and the *tao3*(E1493Q) alleles was selected and crossed with a *trp1-1* (i.e. trp1(D87-stop)) derivative of BY4741. After sporulation of the resulting diploid cell, spores with the following genotype Mat a, LYS2, met15Δ0, ura3Δ0, leu2Δ0, his3Δ1, trp1-1, RME1-Δ, ins-308A, TAO3(E1493Q) or Mat alpha, lys2Δ0, ura3Δ0, leu2Δ0, his3Δ1, trp1-1, RME1-Δ, ins-308A, TAO3(E1493Q) were selected to create strains LMA5395 and LMA5396 respectively. LMA5395 was transformed with plasmid 1692 linearized by *Bsu36I* digestion. A tryptophan prototroph transformant was selected to create strain LMA5419.

*E. coli* strain NEB 10-beta (NEB Cat# C3019) was used for construction of plasmids by ligation-less Gibson assembly (Fu *et al*, 2014) and their multiplication. All the plasmid inserts were verified by sequencing.

C-terminal TAP-tagged strains originated from the collection of systematically built strains (Ghaemmaghami *et al*, 2003). Deletion strains were part of the systematic yeast gene deletion collection (Giaever *et al*, 2002) distributed by EuroScarf (http://www.euroscarf.de) or were built by transformation of BY4741 strain with a cassette containing a selection marker cassette flanked by long recombination arms located upstream and downstream the open reading frame of the gene. Deletions were tested by PCR amplification of the modified locus.

To create auxin-inducible degron strains, the indicated genes were C-terminally tagged with a polyG-3Flag-miniAID-kanMX6 cassette amplified by PCR using plasmid p1451 as a template (Dehecq *et al*, 2018) in LMA5396. The resulting strains were then crossed with LMA5419 and sporulated to recover strains of the desired genotype. The only exceptions were the CCR4-degron and HYP2-degron strains for which the AID-tag amplified from plasmid p1602 has been inserted at the N-terminus of the protein by CRISPR/CAS9 engineering as described (Mans *et al*, 2015).

### Media and growth conditions

Yeast cells were grown in YPD (20g.L^-1^ glucose, 10g.L^-1^ yeast extract, 20g.L^-1^ bacto-peptone, 20g.L^-1^ bactoagar for plates only) and in synthetic media without uracil to select transformants and maintain plasmids with the URA3 marker. All yeast strains were freshly thawed from frozen stocks and grown at 30°C. Bacterial strains were grown in LB media, supplemented with antibiotics when necessary, at 37°C.

### Plasmids construction

#### Plasmids for the His3 codon-changed reporters

Coding sequences for His3 versions, « 10 % » and « 100 % » were recovered from published data (Radhakrishnan *et al*, 2016). The DNA fragments were synthesised by Twist Bioscience (San Francisco, CA, USA). The coding sequences were amplified with LA070 and LA071 before a Gibson assembly reaction. The final PCR product was cloned into the *BamH*I and *Not*I sites of pCM189 (Garí *et al*, 1997).

#### Plasmids for the degron system

For the p1603 plasmid, the Z3EV artificial transcription factor was PCR amplified from plasmid pFS461(Ohira *et al*, 2017) using oligonucleotides FF3765 and FF3766 and the PCR product was used to transform a URA3 derivative of BY4741. 5-FOA resistant cells were selected and checked for correct replacement of the URA3 coding sequence by the Z3EV artificial transcription factor coding sequence. The *ura3::Z3EV* allele obtained was amplified from yeast genomic DNA using oligonucleotides FF293 and FF3746. The final PCR product was digested with *Sst*II and cloned into the *Sst*II and *SmaI* sites of pRS304.

For the p1692 plasmid, the Z3EV promoter was PCR amplified from plasmid pFS478 (Ohira *et al*, 2017) using oligonucleotides FF3740 and FF3872. The obtained PCR product was digested with *SalI* and *BamHI*. The *O. sativa* TIR1 ORF followed by the Nrd1 terminator was PCR amplified using oligonucleotides FF3718 and FF3719 from the genomic DNA of a *nrd1-AID* strain kindly provided by D. Challal (Domenico Libri laboratory, IJM, Paris, France). The PCR product was digested with *BamHI* and *XhoI*. Both PCR products were cloned together into the *XhoI* site of plasmid p1603.

### Cell culture, RNA extraction and degradation kinetics

For RT-qPCR and RNA sequencing, cells were first grown in YPD to log phase and collected. Total RNA was extracted using the hot phenol extraction method and precipitated using ammonium acetate and ethanol.

For the tetOFF promoter inhibition and steady state analysis, cells expressing the appropriate plasmids were grown at 25°C in synthetic media without uracil to allow expression of the reporter mRNA. For the analysis of degron mutants, when cells reached a OD600 of 0.4, β-estradiol for Tir1 expression at a final concentration of 1 μM and IAA auxin at a final concentration of 100 μM for protein depletion were added directly to the media. Several time points were collected to verify protein depletion by western blot with an anti-FLAG HRP. After 1h depletion, cell were harvested for steady state analysis. For transcriptional repression, doxyxycline was added at a final concentration of 10 μg/mL. Cells were collected at the time points indicated in the Northern Blot, generally, 0, 2, 4, 8, 10, 20, 30 minutes for the non-OPT-HIS3 RNA and 0, 5, 10, 20, 30, 60 for the more stable, OPT-HIS3 RNA.

For translation arrest, cycloheximide was added at a final concentration of 5 to 50 μg/mL together with the doxycycline.

### Northern blots

RNAs were separated on 1.5% agarose gels, transferred on nylon membrane (Hybond N+, Amersham, GE) that was UV cross-linked at 0.120 Joules and probed with DIG-labelled RNA or DNA probes. Digoxigenin-containing RNA probes were generated by in vitro transcription with T7 polymerase using the DIG RNA Labelling kit (SP6/T7) from Roche (cat. No 11175025910). The oligonucleotide AJ529 (T7 promoter) pre-annealed with a target-specific oligonucleotide composed of the T7 promoter reverse complement fused to a template sequence, LA94 for His3-tCyc and LA95 for SCR1 were used as a template. Specific DNA probes for the non-optimal and optimal His3 RNA were generated using MFR917 and LA084. After PCR amplification and purification, a single stranded amplification was done using MFR918 for the OPT and the non-OPT version of the probes.

For poly(A) tail visualization, RNAs were cleaved using RNAse H (New England Biolabs, Ipswich, MA, USA) with LA121 as a specific primer for HIS3-tCYC1. Oligo-dT were added to the reaction to remove poly(A) tails for the «A0» reference position. RNAs were then purified with acid phenol-chloroform extraction. Cleaved RNAs were separated on a 6% acrylamide gel, transferred to a nylon membrane (Hybond N+, Amersham) and UV cross-linked. For the DNA ladder, a denatured PBR322 DNA digested with *MspI* was used. Prior to the assay, digoxigenin-11-ddUTP was added to the 3’ end of the digested fragments using terminal transferase enzyme (New England Biolabs, Ipswich, MA, USA).

### RNA half life measurements

A mix of reverse qPCR oligonucleotides, LA126 (Act1), LA102 (His3-tCyc) and random hexamers were used for the reverse transcription. Serial dilutions of cDNA were quantified by TaqMan probe-based quantitative PCR. The amplification was done with the Bio-Rad CFX96 machine and the corresponding software (CFX Maestro Software), with step 1 (95°C for 30 seconds) and step 2 (40 cycles of 95°C for 15 seconds and 60°C for 30 seconds). Probes were LA122-5’-Cy5/TAO/3’IBRQ for His3-tCyc and LA125-5’-HEX/ZEN/3’IBFQ for Act1. Primers were LA102 and LA123 for HIS3, LA124 and LA126 for ACT1.

### Affinity purification for RNA sequencing, immunoblot and mass-spectrometry

TAP-tagged proteins were purified using a one-step purification method(details in Namane & Saveanu, 2022). Frozen cell pellets of 4 L culture were resuspended in lysis buffer (20 mM HEPES K pH 7.4, 100 mM KOAc, 0.5% Triton X100, 5mM MgCl2, Protease inhibitor, 1X Vanadyl-Ribonucleoside Complex, 40 units ml^-1^ RNasin) and lysed using a MagNA Lyser (two passages of 90s at 4000 rpm). The lysate was cleared at 4°C for 20 min at 14000 rpm. Magnetic beads (Dynabeads M-270 epoxy) coupled to IgG were added to the protein extract and incubated for 1,5 h at 4°C (Oeffinger *et al*, 2007). Beads were magnetically separated and extensively washed five times with a washing buffer (20 mM HEPES K pH 7.4, 100 mM KOAc 0.5% Triton X100, 5mM MgCl2). After washing, proteins were eluted by incubation in denaturing buffer (SDS 2% in Tris-EDTA pH 7.5) 10 minutes at 65°C. After collection, RNA were extracted by hot acid phenol/chloroform method, precipitated with ammonium acetate and ethanol. Samples were treated by RiboZero (Illumina) or Ribominus (Thermo Scientific) rRNA removal reagents and libraries were prepared using TruSeq Stranded mRNA kit (Illumina). We strictly followed the protocol except that we started directly at the fragmentation step after the rRNA removal step and we adjusted the number of cycle required to amplify libraries. For RNAse treatment, 1 ul of micrococcal nuclease (Biolabs M02475, 2×10^6^ U/ml) was added to 200 μl buffer containing 1 mM CaCl_2_ and incubated for 10 min 37°C. An RNA aliquot was harvested prior to TAP purification for the analysis of total RNAs (input) and was treated the same way as the purified fractions except for the purification.

### Illumina RNA sequencing results analysis

Reads were aligned along the 16 chromosomes and mitochondrial sequence of *S. cerevisiae* S288C genome. For the mapping, we used the STAR program (version 2.7) with default parameters except for the following preferences: alignIntronMax 1500, alignMatesGapMax 1500, alignSJoverhangMin 25. For the exon annotation we used the GTF file for *Saccharomyces cerevisiae* (R64-1-1.104) from ENSEMBL. Read counts were obtained using the featureCounts function of the Subread package, with an annotation file that contains non-coding RNA coordinates in addition to yeast gene transcripts (Malabat *et al*, 2015). The analyzed data was visualized using the Integrated Genome Viewer, IGV (Robinson *et al*, 2011) and further data processing was done using R (R Core Team, 2022). It consisted in filtering out features with less than 10 reads, followed by a normalization by the total number of reads for each sample and averaging of the log2 transformed results for the three replicated experiments for each condition. Input normalized counts were substracted from purification counts for each condition. Finally, the results were adjusted by substracting the median of the log2 enrichment values for each quantified feature. For the comparison with RT-qPCR results, raw enrichment values were calculated by substracting, for each replicate, the fraction of counts in input samples from the fraction of counts in each purified sample (log2 transformed values).

### Reverse transcription and quantitative PCR

The extracted RNA samples were treated with DNase I (Ambion TURBO DNA-free kit) before reverse-transcription (RT) with the Superscript II or III (Invitrogen) and library preparation. The purified and input samples were used for reverse transcription with transcript-specific primers. A mix of reverse qPCR oligonucleotides, CS888 (RPL28-premature), CS889 (RPL28) and random hexamers were used for the RT. Serial dilutions of cDNA were quantified by qPCR, CS887-CS888 (RPL28-premature) and CS889-CS946 (RPL28). The amplification was done in a Stratagene MX3005P and corresponding software (MXpro quantitative PCR system), with step 1 (95°C for 5 min) and step 2 (40 cycles of 95°C for 20 s and 60°C for 1 min).

Half-life estimates from RT-qPCR results were done using nonlinear regression on values that were calculated as a fraction from RNA levels at time 0, using the exponential decay function e^-(t-l)^*^k^. In this fit, “*t*” is time after doxycyclin addition, “*l*” is a lag period (Baudrimont *et al*, 2017), estimated from the results obtained with the wild-type condition, and *“k”* is the decay constant. Estimates of half-life were obtained by the formula (ln(2)/*k*)+*l*, that takes into account the lag period. Initial estimates of this lag, based on both non-OPT HIS3 and OPT HIS3 reporters, led to a value of 1.2 min. that was used throughout to obtain estimates. The *confint* function from the “MASS” R package was used to obtain confidence estimates for “*k*” and associated half-life values at a 0.95 level of confidence.

### Protein extracts and immunoblots

Total protein extracts were prepared from 5 ml of exponential culture cells using an alkaline treatment. Cells were incubated in 200 μL of 0.1M NaOH for 5 min at room temperature, collected by 3 min centrifugation and resuspended in 50 μL of SDS sample buffer containing DTT (0.1M). Proteins were denatured for 3 min at 95°C, and cellular debris were pelleted by centrifugation. 10 μL of supernatant or diluted supernatant (for quantification scale) were loaded on acrylamide NuPAGE Novex 4-12% Bis-Tris gels (Life technologies). Transfer to a nitrocellulose membrane was done with a semi-dry fast system (Biorad trans-blot) with discontinuous buffer (BioRad technote 2134). Proteins were detected by hybridization with anti-FLAG-HRP, for the detection of the FLAG tag (A8592, Sigma-Aldrich, clone M2, monoclonal), PAP (P1291, Sigma-Aldrich, peroxidase anti-peroxidase soluble complex antibody), for the detection of the protein A fragment of the TAP tag or anti-HA peroxidase 1/500 (clone 3F10, Roche).

### Mass spectrometry acquisition and data analysis

Protein samples were treated with Endoprotease Lys-C (Fujifilm Wako chemicals, Osaka, Japan) and Trypsin (Trypsin Gold Mass Spec Grade, Promega). Peptide samples were desalted using OMIX C18 pipette tips (Agilent Technologies). The peptides mixtures were analyzed by nano-LC-MS/MS using an Ultimate 3000 system (Thermo Fisher Scientific) coupled to an LTQ-Orbitrap Velos mass spectrometer. Peptides were desalted on-line using a trap column (C18 Pepmap100, 5μm, 300μmÅ∼5mm (Thermo Scientific) and then separated using 120min RP gradient (5−45% acetonitrile/0.1% formic acid) on an Acclaim PepMap100 analytical column (C18, 3μm, 100., 75μm id x 150mm, (Thermo Scientific) with a flow rate of 0.340μL.min^-1^. The mass spectrometer was operated in standard data dependent acquisition mode controlled by Xcalibur 2.2. The instrument was operated with a cycle of one MS (in the Orbitrap) acquired at a resolution of 60,000 at m/z 400, with the top 20 most abundant multiply-charged (2+ and higher) ions subjected to CID fragmentation in the linear ion trap. An FTMS target value of 1e6 and an ion trap MSn target value of 10000 were used. Dynamic exclusion was enabled with a repeat duration of 30s with an exclusion list of 500 and exclusion duration of 60s. Lock mass of 445.12002 was enabled for all experiments. The results were analyzed with MaxQuant and processed with Perseus and R, as previously described (Dehecq *et al*, 2018). Enrichment calculations were based on a standard data set of protein abundance in yeast (Ho *et al*, 2018). At least three independent experiments were performed for each purified protein.

### Nanopore sequencing and analysis

Degradation of target proteins was induced for 1 hour in exponential growth cultures in rich (YPD) medium, with β-estradiol (1μM final) and IAA (100μM final). Protein depletion was verified by immunoblot against the FLAG tag. Total RNA was extracted using glass beads and hot phenol protocols. RNA libraries were prepared from 250 to 500 ng of oligo-(dT)25-enriched mRNA from with a Direct RNA Sequencing Kit (catalog no. SQK-RNA002, Oxford Nanopore Technologies) according to the manufacturer’s instructions. Sequencing was performed using R9.4 flow cells on a MinION device (ONT). Raw data were basecalled using Guppy (ONT). Reads were mapped to the SC288 reference genome Saccer3.fa using minimap2.1 with options “-k 14 -ax map-ont -secondary = no” and processed with samtools 1.9 (samtools view -b -o). The poly(A) tail lengths for each read were estimated using the Nanopolish 0.13.2 polya function (Workman *et al*, 2019). In subsequent analyses, only length estimates with the QC tag that was reported by Nanopolish as “PASS” were considered. We also retained only features with at least 20 assigned reads in two independent replicates. Statistics of obtained results, poly(A) tail length and counts, were computed with R.

### Simulation of RNA degradation and deadenylation

To be able to estimate how global poly(A) tail would change with RNA stability, we used two different approaches. In one, we obtained, using Maxima (Maxima, 2022), an ordinary differential equation solving system, the equations describing the evolution over time of three species of RNA of different poly(A) length (long, medium and short) under two kinetic models. The deadenylation-dependent, or serial, model described by the following expressions:

ser1:’diff(PAl(t), t)=T-kA*PAl(t);

ser2:’diff(PAm(t), t)=kA*PAl(t)-kA*PAm(t);

ser3:’diff(PAs(t), t)=kA*PAm(t)-kD*PAs(t).

Here, “T” is constant and represents all the events preceding RNA deadenylation and degradation (synthesis, splicing, nuclear export), “kA” is a deadenylation constant (pseudo-first order process) that dictates the rate of transformation of long to medium and to short poly(A) mRNAs and “kD” is a degradation constant, corresponding to the transformation of the RNA to decay products. “t” represents time. Solving the system involves the following Maxima commands:

atvalue(PAl(t), t=0, 0); atvalue(PAm(t), t=0, 0); atvalue(PAs(t), t=0, 0);

desolve([ser1, ser2, ser3], [PAl(t), PAm(t), PAs(t)]);

For the deadenylation-independent, or parallel, model, the equations included an additional step of degradation for both PAl and PAm species:

par1:’diff(PX1(t), t)=T-kA*PAl(t)-kD*PAl(t);

par2:’diff(PX2(t), t)=kA*PAl(t)-kA*PAm(t)-kD*PAm(t);

par3:’diff(PAs(t), t)=kA*PAm(t) - kD*PAs(t);

atvalue(PAl(t), t=0, 0); atvalue(PAm(t), t=0, 0); atvalue(PAs(t), t=0, 0);

desolve([par1, par2, par3], [PAl(t), PAm(t), PAs(t)]);

For steady-state conditions, with “t” reaching very high values, the obtained results indicated that PAl and PAm accumulation would be only dependent on deadenylation, k_A_, rates in the deadenylation-dependent model. PAs accumulation at steady state would be only dependent on k_D_, as expected. For the deadenylation-independent model, the relative levels of the different species of RNA at steady state depend on both k_A_ and k_D_.

While the obtained equations could have been used directly, we prefered to perform an independent validation using the Tellurium system for modeling (Choi *et al*, 2018), which also allows flexibility in the choice of initial conditions and parameters. Steady-state values obtained from the formulas obtained with Maxima were fed into a Tellurium model that can be described by the following expressions:

model rnadeg_deadenylation_dependent

$T -> PAl; kS*T

PAl -> PAm; kA*PAl

PAm -> PAs; kA*PAm

PAs -> ; kD*PAs

and

model rnadeg_deadenylation_independent

$T -> PAl; kS*T

PAl -> PAm; kA*PAl

PAl -> ; kD*PAl

PAm -> PAs; kA*PAm

PAm -> ; kD*PAm

PAs -> ; kD*PAs

Variations in k_D_ and k_A_ values were used to follow the change in the simulated species amounts of RNA over time, in the absence of new RNA generation (models missing the generation of PAl). The sum of the three species and its decrease over time were analyzed similar to how real experimental data are processed to obtain estimates of half-life. The PAl, PAm and PAs were assigned arbitrary 10 values of 70, 40 and 10 nucleotides, to be able to calculate average simulated poly(A) tail length over time.

### Strains

Strains:
| Strain | Genotype | Reference |
| --- | --- | --- |
| LMA2154 (BY4741) | MATa ura3Δ0 his3Δ1 leu2Δ0 met15Δ0 | Brachmann et al. *Yeast*, 1998 |
| LMA5395 | TAO3_E1493Q/RME1(YGR044c) insert A -308/trp1(YDR007w)_D87Stop (trp1-1) | This study, Bushkin et al., *Nat Commun* 2019 |
| LMA5396 | TAO3_E1493Q/RME1(YGR044c) insert A -308/trp1(YDR007w)_D87Stop (trp1-1) | This study, Bushkin et al., *Nat Commun* 2019 |
| LMA5419 | TAO3_E1493Q/RME1(YGR044c) insert A -308/trp1-1::TRP1-prURA3-Z3EVtf-tURA3-Z3EV-TIR1 | This study, Bushkin et al., *Nat Commun* 2019 |
| LMA5575 | LMA5419 + DCP2(YNL118c)::polyG-3Flag-AID::KANMX | This study, inducible degron strain DCP2 |
| LMA5593 | LMA5419 + CCR4(YAL021C)::polyG-3Flag-AID::KANMX6 | This study, inducible degron strain CCR4 |
| LMA5597 | LMA5419 + POP2(YNR052c)::AID-KANMX6 | This study, inducible degron strain POP2(CAF1) |
| LMA5601 | LMA 5419 AID-3Flag-CCR4/POP2(YNR052c)::AID-KanMX6 | This study, degron strain POP2(CAF1)/CCR4 |
| LMA2194 | BY4741 UPF1-TAP::HIS3MX6 | Ghaemmaghami et al., *Nature* 2003 |
| LMA2195 | BY4741 RPL16A-TAP::HIS3MX6 | Ghaemmaghami et al., *Nature* 2003 |
| LMA4987 | BY4741 PAB1-TAP::HIS3MX6 | Ghaemmaghami et al., *Nature* 2003 |
| LMA4985 | BY4741 LSM1-TAP::HIS3MX6 | Ghaemmaghami et al., *Nature* 2003 |
| LMA4986 | BY4741 LSM7-TAP::HIS3MX6 | Ghaemmaghami et al., *Nature* 2003 |
| LMA2356 | BY4741 PAT1-TAP::HIS3MX6 | Ghaemmaghami et al., *Nature* 2003 |
| LMA5243 | BY4741 DHH1-TAP::HIS3MX6 | Ghaemmaghami et al., *Nature* 2003 |
| LMA1667 | BY4741 *upf1*Δ ::KANMX6 | Giaever et al., *Nature*, 2002 |
| LMA5018 | BY4741 PAB1-HA::KANMX6 | This study |
| LMA5028 | BY4741 PAB1-HA::KANMX6 UPF1-TAP::HIS3MX6 | This study, Ghaemmaghami et al., *Nature* 2003 |

### Oligonucleotides

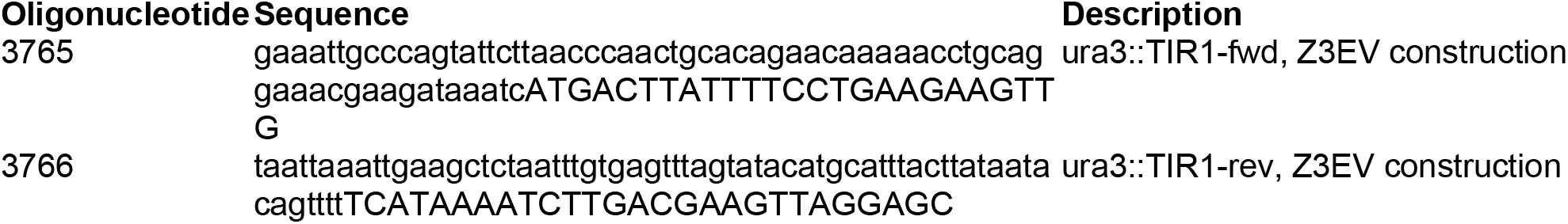
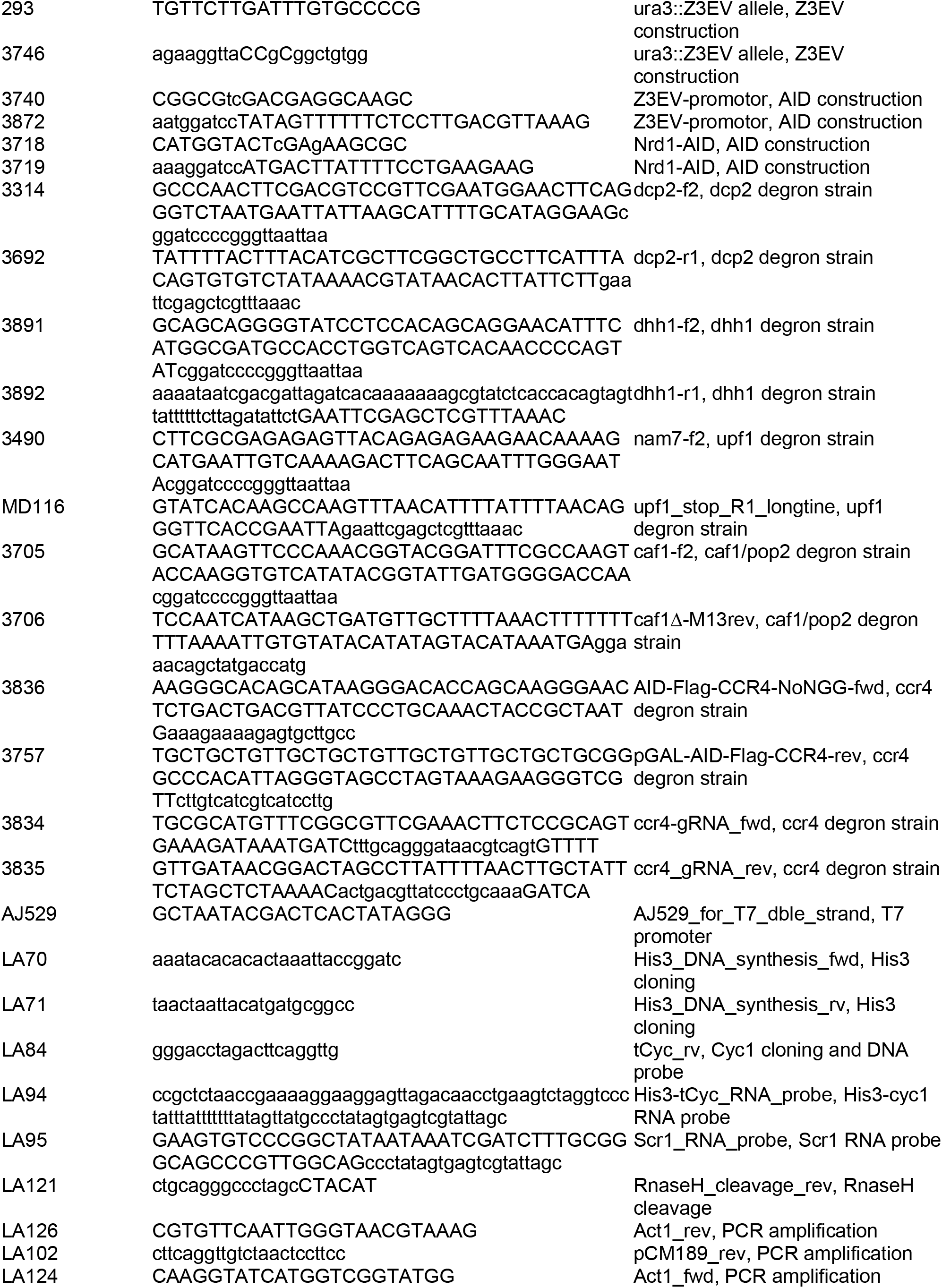
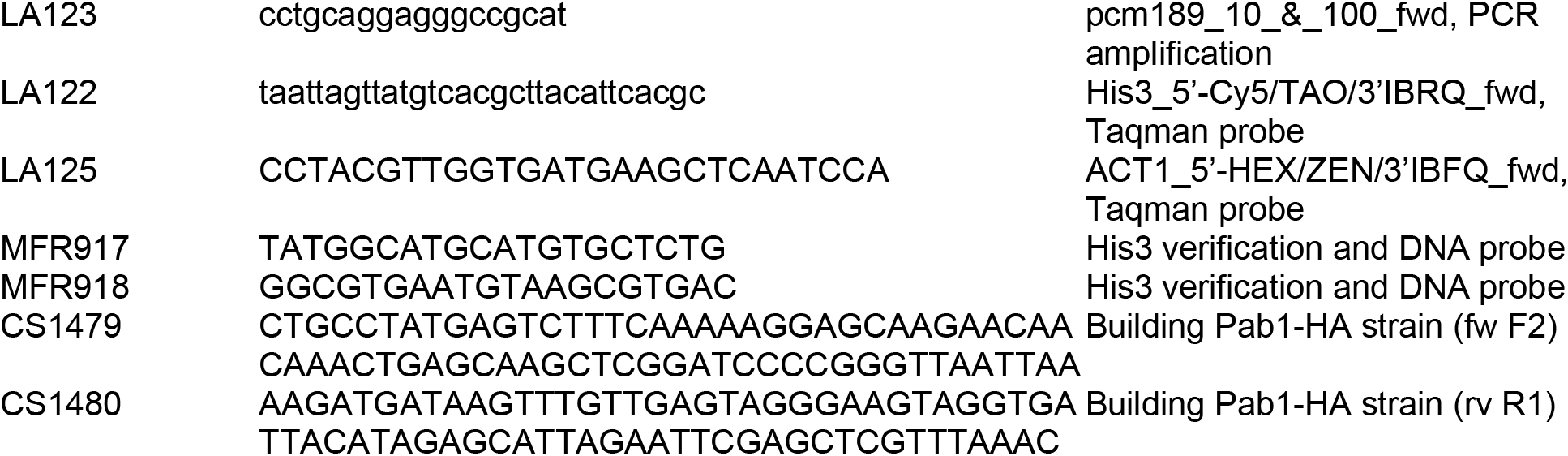

### Plasmids

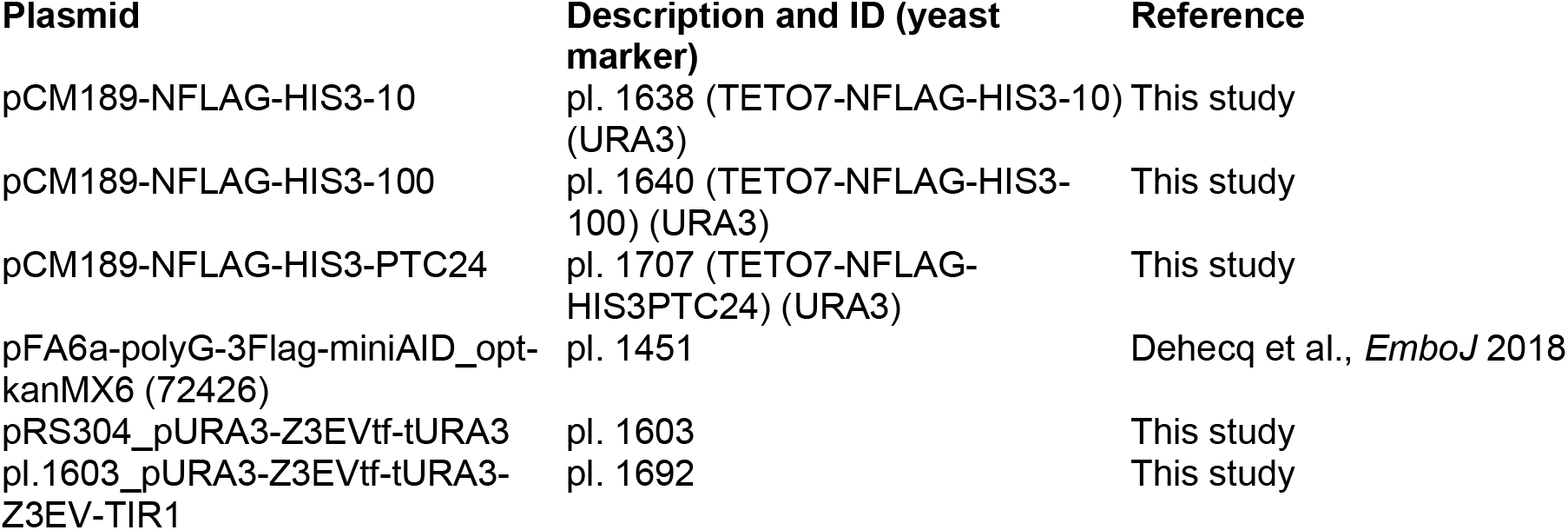

**Plasmids:**
| Plasmid | Description and ID (yeast marker) | Reference |
| --- | --- | --- |
| pCM189-NFLAG-HIS3-10 | pl. 1638 (TETO7-NFLAG-HIS3-10) (URA3) | This study |
| pCM189-NFLAG-HIS3-100 | pl. 1640 (TETO7-NFLAG-HIS3-100) (URA3) | This study |
| pCM189-NFLAG-HIS3-PTC24 | pl. 1707 (TETO7-NFLAG-HIS3PTC24) (URA3) | This study |
| pFA6a-polyG-3Flag-miniAID_opt-kanMX6 (72426) | pl. 1451 | Dehecq et al., *EmboJ* 2018 |
| pRS304_pURA3-Z3EVtf-tURA3 | pl. 1603 | This study |
| pl.1603_pURA3-Z3EVtf-tURA3-Z3EV-TIR1 | pl. 1692 | This study |

## Supplementary Figures

**Fig. S1.**
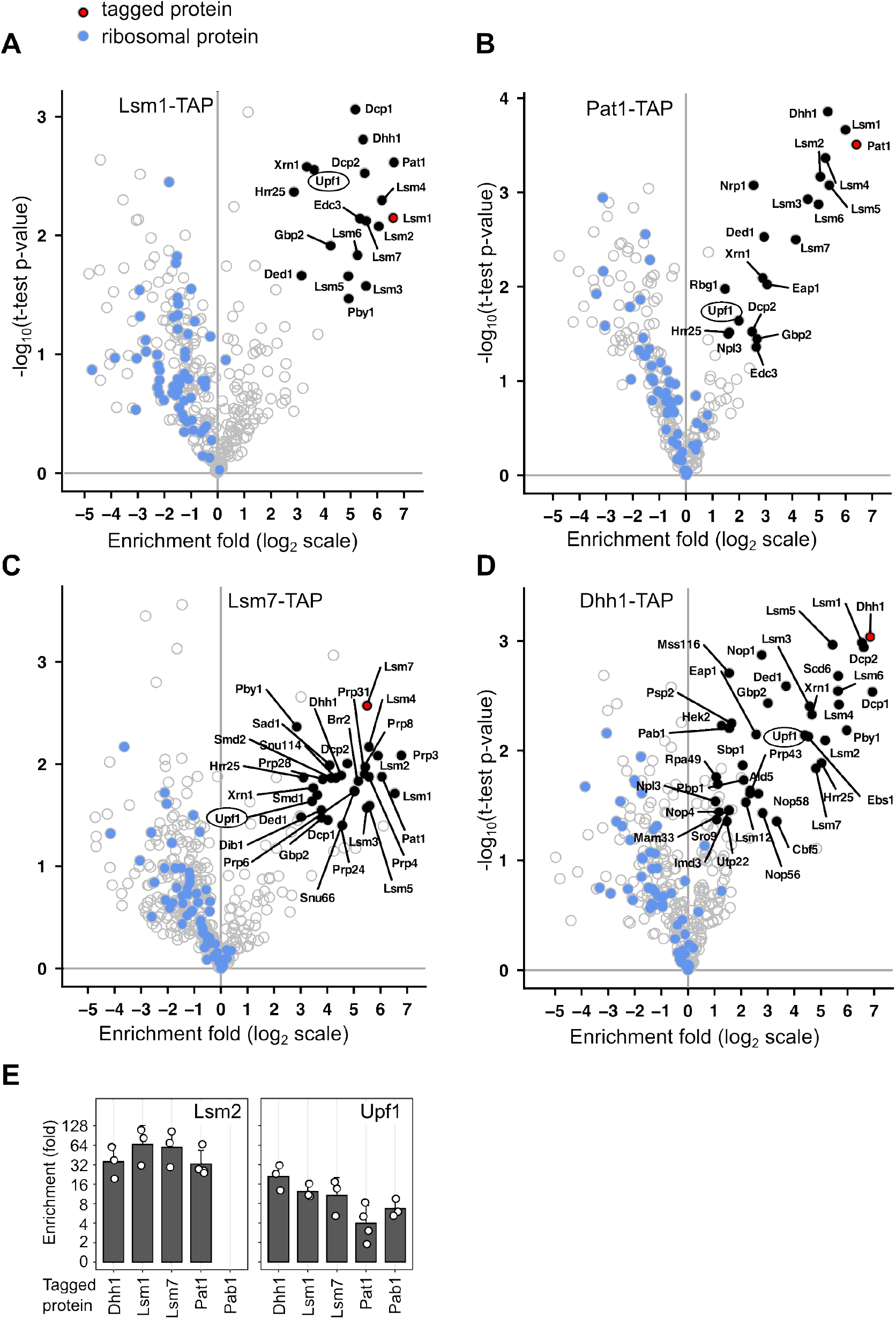
Enrichment of specific factors in purified complexes containing Upf1. Volcano plots represent the enrichment of proteins, relative to the protein abundance in a total extract as measured by label-free mass spectrometry. Results are presented for purifications using TAP tagged versions of Lsm1 **(A)**, Pat1 **(B)**, Lsm7 **(C)**, and Dhh1 **(D)**. Ribosomal proteins are highlighted in blue. **(E)** Examples of enrichment values across the various purifications for Lsm2, a component of Lsm1-7 and Lsm2-8 complexes, and Upf1.

**Fig. S2.**
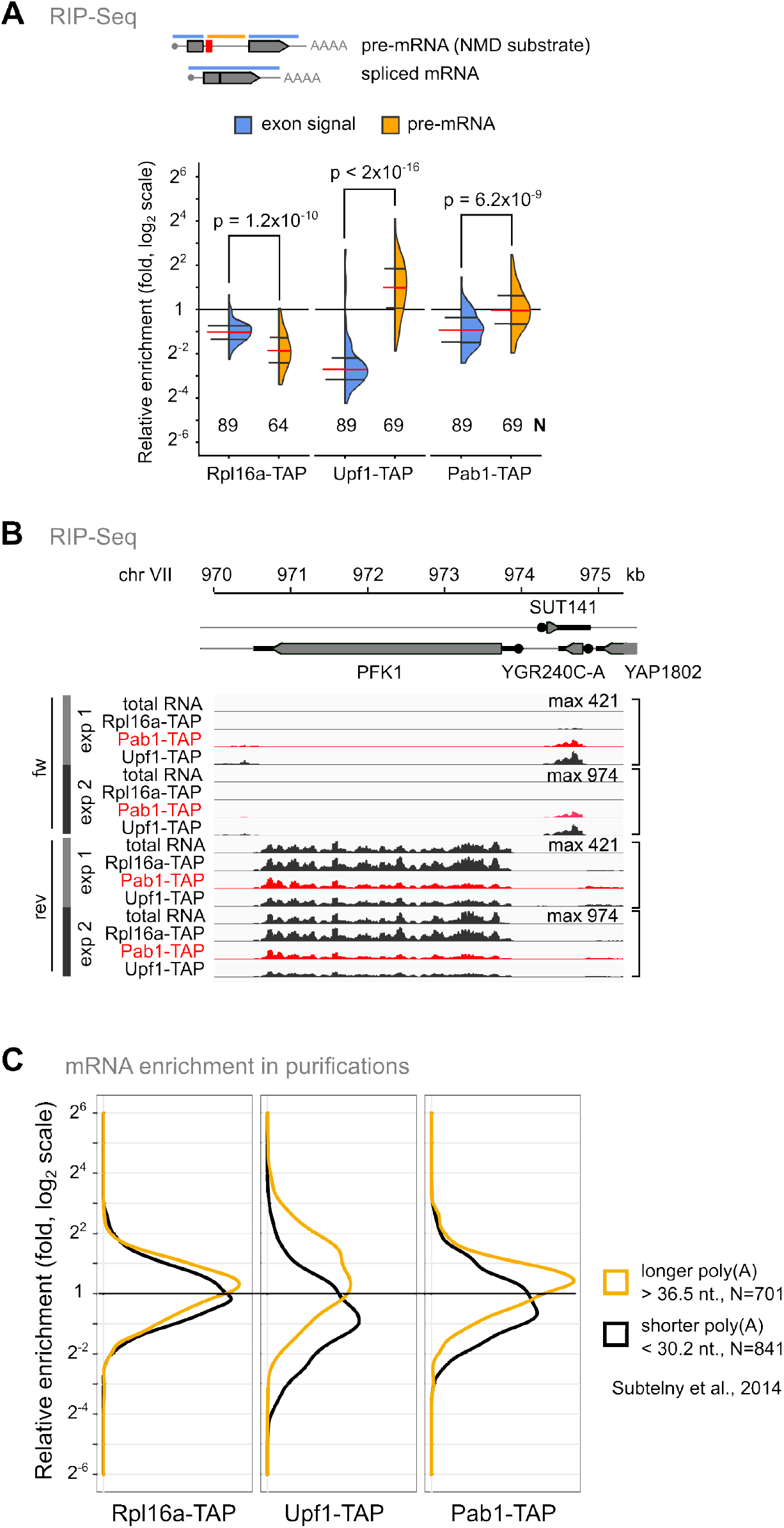
Unstable mRNAs are associated with Pab1. **(A)** Relative enrichment of pre-mRNA for ribosomal protein genes (intron signal, orange) in comparison with the spliced mRNA for the same category (exon signal, blue). The red horizontal line indicates the median of the enrichment values, with the first and third quartiles indicated. The p-values correspond to a Wilcoxon rank sum test with continuity correction, N is the number of values in each category. **(B)** Example of signal intensity for RNAs associated with Pab1-TAP, Rpl16a-TAP or Upf1-TAP for a region of the yeast genome that corresponds to SUT141, an unstable RNA, in comparison with the PFK1 stable mRNA (transcribed in proximity but from the opposite strand). The vertical scale is the same for the samples of the same replicated experiment. Pab1 associated RNA signal is depicted in red. The image was obtained with the Integrated Genome Viewer, IGV. **(C)** The distribution of the enrichment values for two extreme categories of mRNA, with long (orange) or short average poly(A) tails (black) was depicted for the purified samples in association with Rpl16a-TAP, Upf1-TAP and Pab1-TAP.

**Fig. S3.**
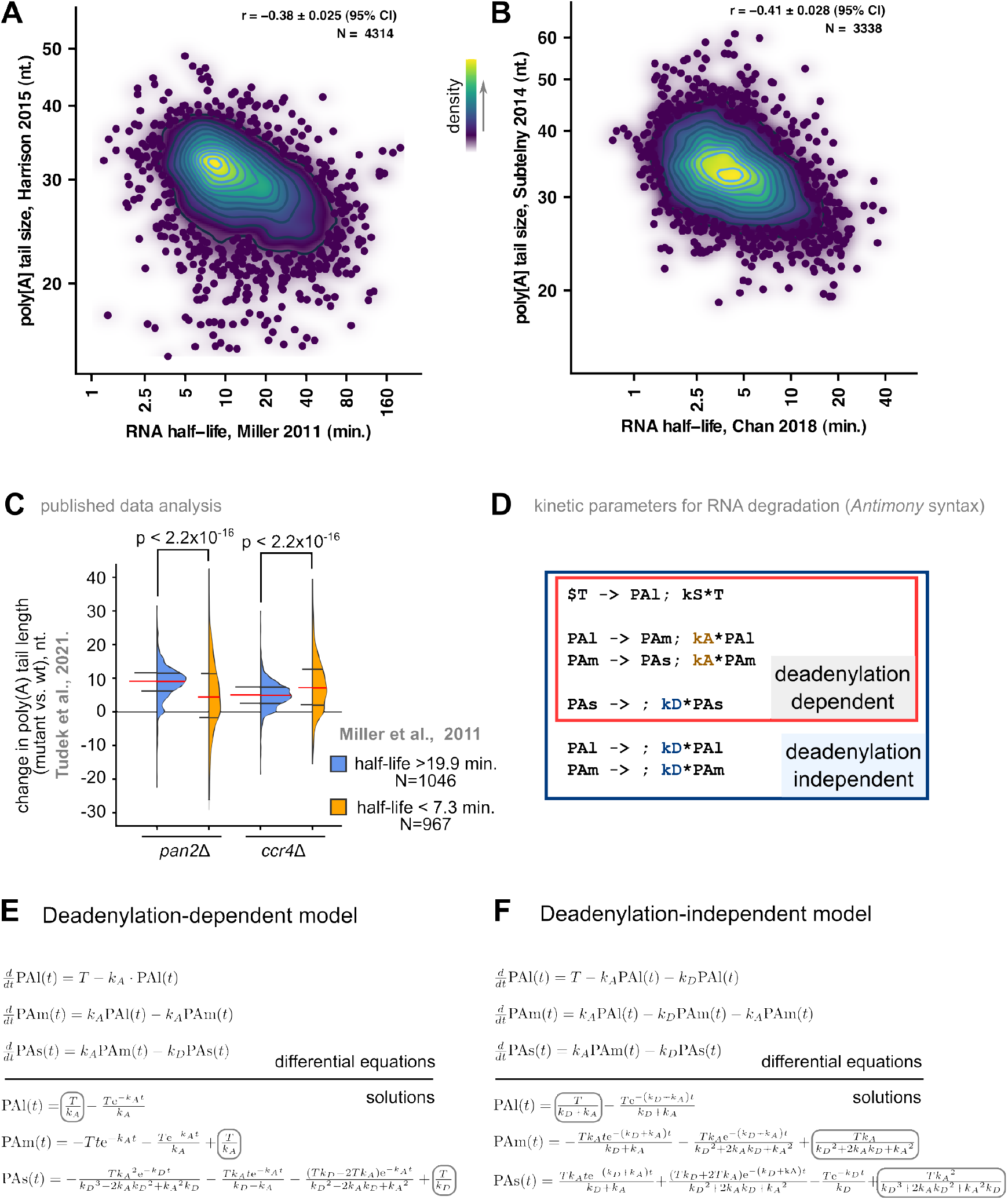
Robustness of the negative correlations between poly(A) tail length and half-life of RNA. **(A), (B)** Negative correlations between published results, similar to the ones shown in Fig. 2A and 2B, including results for poly(A) tail size (Harrison *et al*, 2015; Subtelny *et al*, 2014) and estimates of RNA half-life (Chan *et al*, 2018; Miller *et al*, 2011). **(C)** RNAs for which half life is short, in the first quartile, half life inferior to 7.3 min, or long, in the third quartile, over 19.9 min (Miller *et al*, 2011), were selected to evaluate how their poly(A) size changed in deadenylation mutants, as measured by Nanopore sequencing (Tudek *et al*, 2021). The quartiles of the distributions are indicated, median in red, as well as the p-value of a Wilcoxon rank sum test with continuity correction for the difference between the two populations. **(D)** Parameters used to simulate RNA degradation measurements that include poly(A) tail information and kinetic equations using the *Antimony* syntax (Smith *et al*, 2009). “T” represents all the processes leading to mRNA formation. PAl, PAm and PAs represent concentrations of different poly(A) tail RNA species. **(E)** The set of differential equations used to estimate time changes for the three modeled poly(A) species in a deadenylation-dependent model (upper region) and the solutions (lower region). The terms that correspond to steady-state are circled in gray. **(F)** Similar to (E), for the deadenylation-independent model.

**Fig. S4.**
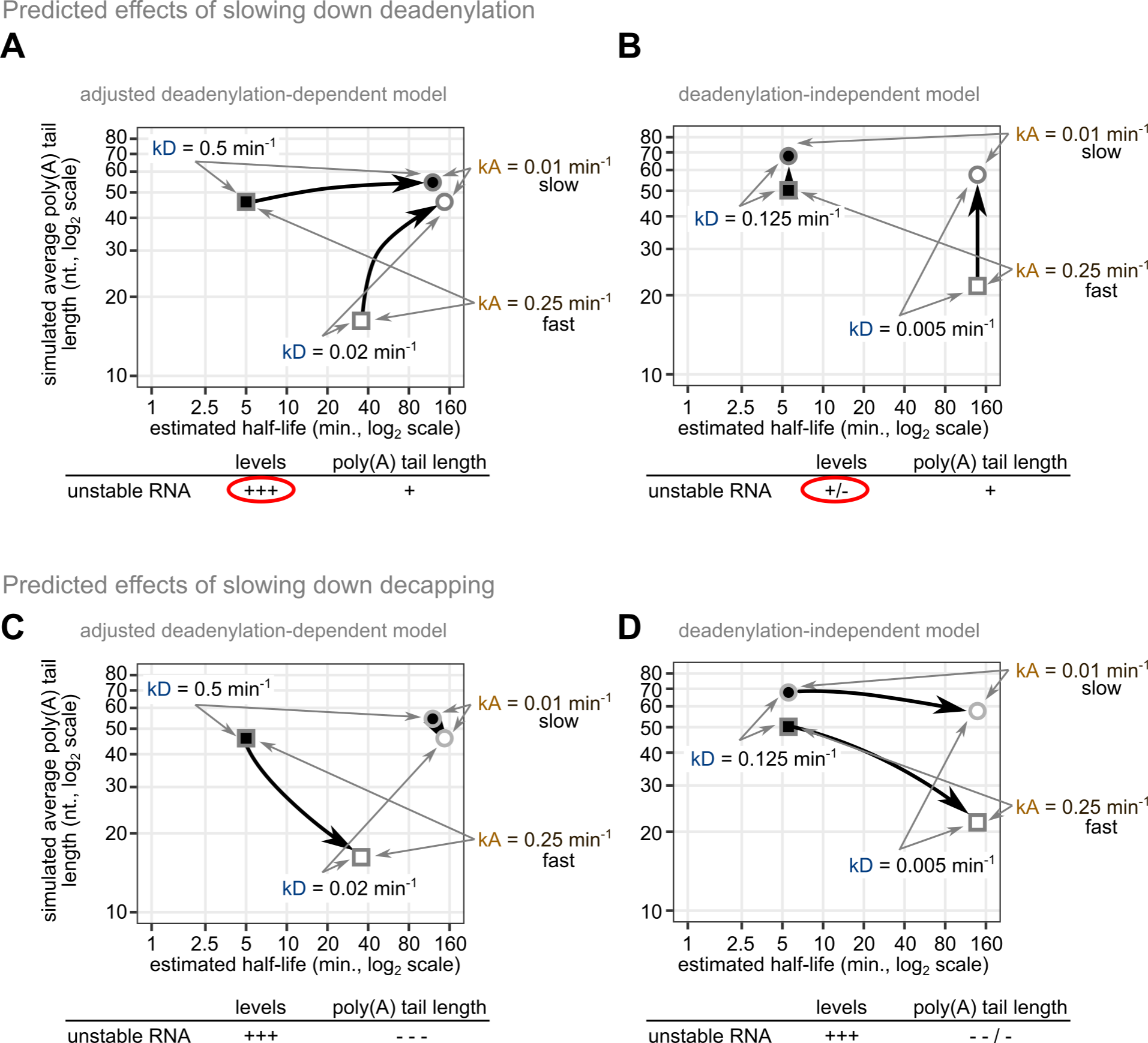
Model-based scenarios for predicted effects of slowing down deadenylation (A, B) and decapping (C, D). For each situation, the expected shift in poly(A) tail length and stability for a stable and an unstable RNA are indicated by black arrows. The experimental conditions can only estimate changes in poly(A) tails and levels for relatively unstable mRNAs (situation depicted under each graph).

**Fig. S5.**
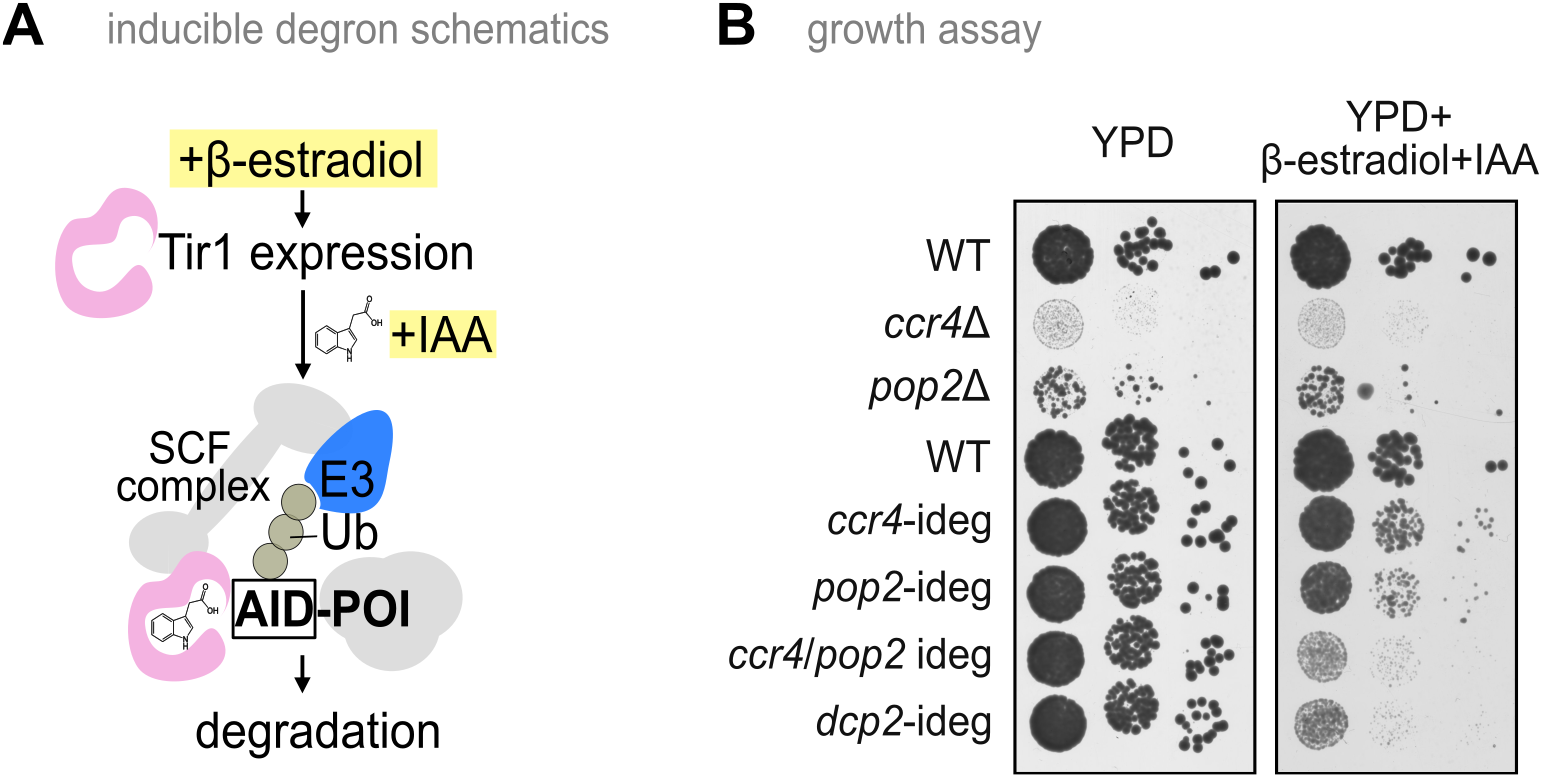
An inducible degron system for deadenylases. **(A)** Schematics of the developed inducible degron system in which *O. sativa* Tir1 expression is under the control of an estrogen sensitive promoter. In the presence of the plant auxin hormone indole-3-acetic acid (IAA), Tir1 targets a E3 ubiquitin ligase to the AID domain fused to the protein of interest (POI) for its rapid proteasome degradation. **(B)** Serial dilution plate growth assay for deletion and inducible degron (ideg) strains affecting deadenylases of the CCR4/NOT complex and the decapping enzyme Dcp2. Growth on rich medium (YPD, left) was compared with growth on medium containing β-estradiol and IAA (right).

**Fig. S6.**
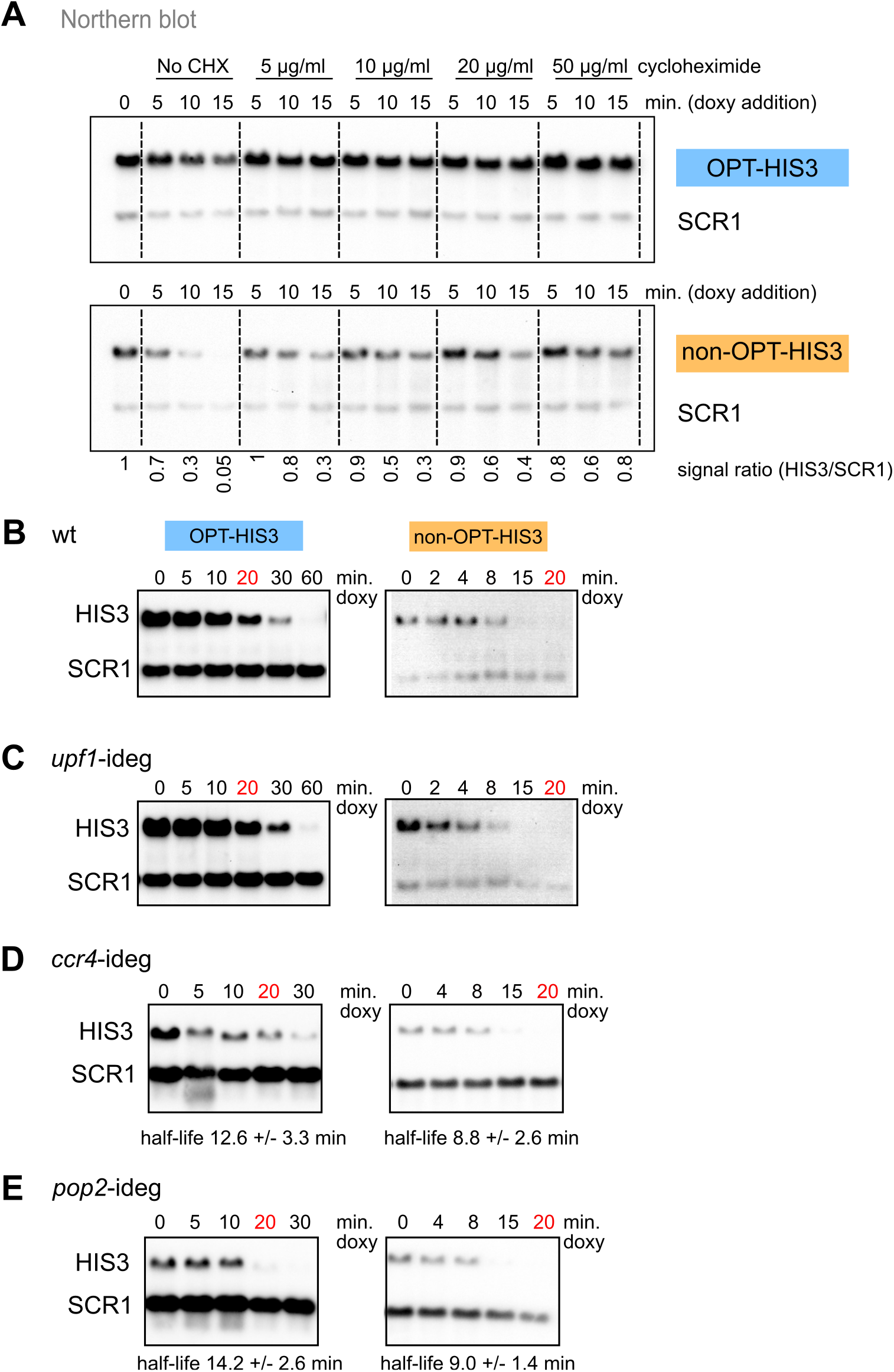
Behavior of OPT-HIS3 and non-OPT-HIS3 reporter mRNA shows that their degradation depends on translation and decapping. **(A)** Dose-dependent effect of the translation inhibitor cycloheximide (from 0 to 50 μg/ml) was tested by northern blotting for 5 optimal (upper panel) and non-optimal (lower panel) HIS3 reporters. SCR1 was used as a loading control. Degradation rates for optimized HIS3 (left panel) and non-optimal HIS3 (right panel) were visualized by northern blot (HIS3 reporter signal) in comparison with an SCR1 control in a wild type strain **(B)** after depletion of Upf1 **(C)**, Ccr4 **(D)**, and Pop2 **(E)**. For each situation, the estimated half -life and the 95% confidence interval obtained from RT-qPCR and the results are presented.

**Fig. S7.**
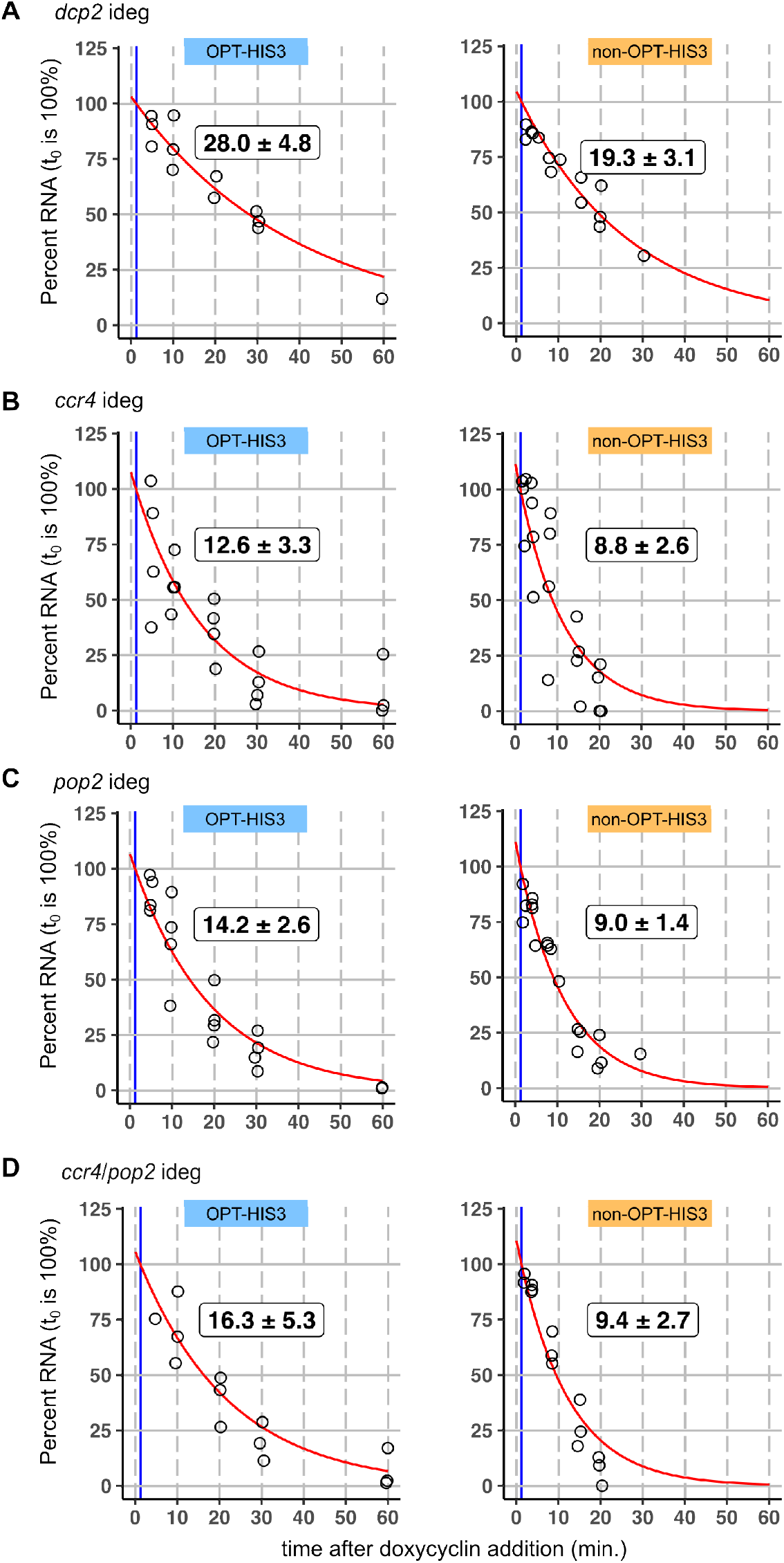
Half-life of reporter RNAs as measured by RT-qPCR. Impact of depletion of Dcp2 **(A)**, Pop2 **(B)**, Ccr4 **(C)**, or the double depletion of Ccr4 and Pop2 **(D)** on the estimated half-life of HIS3 reporter RNA, codon-optimized (left panel) or non-optimal (right panel). Indicated estimates correspond to half-life in minutes with a 95% confidence interval. All the experiments were performed independently at least three times. The wild-type situation is presented in **Fig. 4B**.

**Fig. S8.**
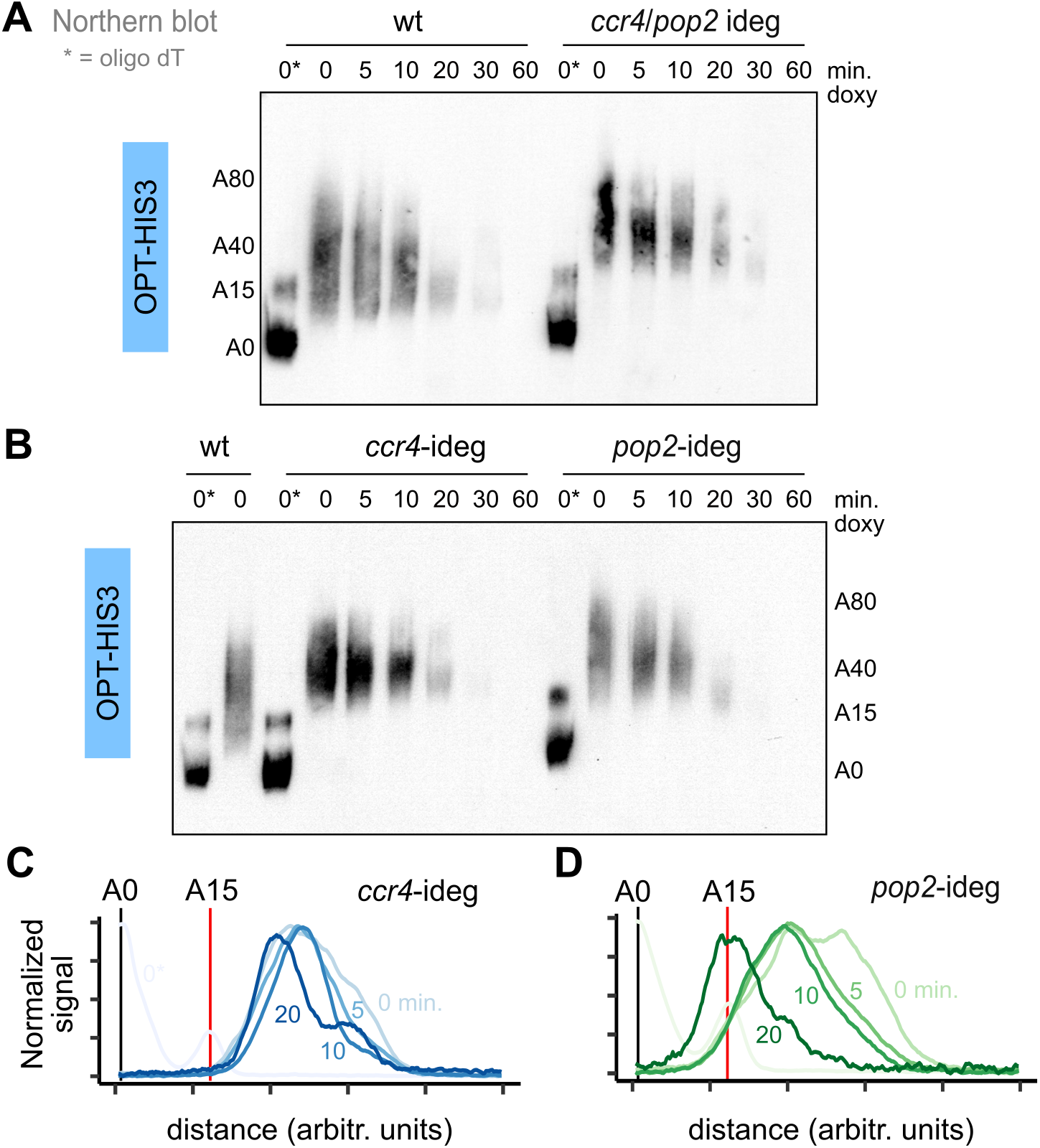
Deadenylation inhibition increases poly(A) tail sizes of reporter RNA, without an effect on their apparent half-life. **(A)** Evolution of poly(A) tail length after transcription shut off with doxycycline in a wild type strain (left side) or after depletion of Ccr4 and Pop2 through the 5 inducible degron system (right side). A star indicates samples treated with oligo-dT prior to RNAse H digestion. Approximate sizes of the poly(A) tails are indicated. **(B)** Similar to (A), but for depletion of Ccr4 or Pop2. A wild type steady-state RNA sample is also shown for comparison with (A). **(C)** and **(D)** Poly(A) tail signal for the experiments presented in (B). The values were normalized to the same maximum signal and are not indicative of the amount of remaining RNA at different time points, only of the distribution. The partially digested signal corresponding to A15 was used to calibrate the plots. The horizontal axis corresponds to distance from the region of the A0 signal.

